# Evolutionary graph theory beyond pairwise interactions: higher-order network motifs shape times to fixation in structured populations

**DOI:** 10.1101/2021.06.26.450017

**Authors:** Yang Ping Kuo, Oana Carja

## Abstract

To design population topologies that can accelerate rates of solution discovery in directed evolution problems or in evolutionary optimization applications, we must first systematically understand how population structure shapes evolutionary outcome. Using the mathematical formalism of evolutionary graph theory, recent studies have shown how to topologically build networks of population interaction that increase probabilities of fixation of beneficial mutations, at the expense, however, of longer fixation times, which can slow down rates of evolution under elevated mutation rate. Here we find that moving beyond dyadic interactions is fundamental to explain the trade-offs between probability and time to fixation. We show that higher-order motifs, and in particular three-node structures, allow tuning of times to fixation, without changes in probabilities of fixation. This gives a near-continuous control over achieving solutions that allow for a wide range of times to fixation. We apply our algorithms and analytic results to two evolutionary optimization problems and show that the rate at which evolving agents learn to navigate their environment can be tuned near continuously by adjusting the higher-order topology of the agent population. We show that the effects of population structure on the rate of evolution critically depend on the optimization landscape and find that decelerators, with longer times to fixation of new mutants, are able to reach the optimal solutions faster than accelerators in complex solution spaces. Our results highlight that no one population topology fits all optimization applications, and we provide analytic and computational tools that allow for the design of networks suitable for each specific task.

## Introduction

Population structure is a powerful determinant of a population’s evolutionary outcome. Some structures have topological properties that amplify the spread of a mutant with the slightest selective advantage (Pavlogiannis et al., 2018), while others work against the force of selection, turning evolution into a process of pure chance (Lieberman et al., 2005). Evolutionary graph theory is the mathematical framework that formalizes the representation of population structure and its evolutionary effects (Lieberman et al., 2005; Débarre et al., 2014; Szabó and Fath, 2007; Kuo et al., 2021). In this framework, each individual occupies a node in a graph, and the edges represent the spatial or replacement patterns of interaction between neighboring nodes. The mode and tempo of evolution are studied through two main quantities: probabilities of fixation, which measure the likelihood that the mutant lineage takes over the whole population, and times to fixation, the expected times until the population consists of only mutant descendants.

Most prior theoretical work has focused on studying the probability of fixation, with theoretical explorations of time to fixation restricted to small networks (Hindersin et al., 2016), symmetric topologies such as lattices, rings, and stars (Paley et al., 2007; Hathcock and Strogatz, 2019; Altrock et al., 2017; Askari and Samani, 2015) or sparse networks, where mutants grow in clusters (Frean et al., 2013; Hajihashemi and Samani, 2019). This imbalance of focus can be partially attributed to the assumption that the waiting time until a successful mutant appears in the population is much larger than the time it takes for this mutant to sweep through it. This would make the rate of evolution in the population mostly depend on the mutant’s probability of fixation. However, this assumption does not always hold, especially for engineering applications with the goal of increasing rates of discovery in directed evolution and evolutionary optimization (Lai et al., 2004; Myers et al., 1985; Bridges and Woodgate, 1985; Greener et al., 1997; McCullum et al., 2010). These applications often utilize an elevated rate of mutation in order to speed up the generation of new candidate solutions, which accelerates search time by several orders of magnitude (Moore and Arnold, 1996; Moore et al., 1997). In these contexts, fixation times of new mutants can shape rates of evolution more than fixation probabilities, and selecting population structures purely for amplification of selection could lead to a substantial evolutionary slowdown (Frean et al., 2013).

This has lead to recent interest in studying times to fixation, especially in the context of trade-offs with the probability of fixation (Möller et al., 2019; Tkadlec et al., 2019). These studies have explored how to topologically build population graphs that increase probabilities of fixation of beneficial mutations, at the expense, however, of longer fixation times, which can slow down rates of evolution under elevated mutation rate. It remains an open question how to change a network’s topology in order to optimize times to fixation of new variants in the population, with negligible change to probabilities of fixation, i.e. to the amplification of selection.

While previous work has focused on pairwise interactions exclusively, many graph connections and spatial patterns of replacement do not take place between pairs of nodes, but rather as collectives at the level of groups of nodes (Alvarez-Rodriguez et al., 2021; Benson et al., 2016). Here we show that moving beyond dyadic structures is fundamental to explain the trade-offs between probability and time to fixation and to be able to understand how to tune network properties to minimize or maximize times to fixation without changing probabilities of fixation. Networks exhibit higher-dimensional patterns of interconnections which can be organized by the number of nodes participating in forming the patterns (**Figure 1A**). Lower-order patterns of connectivity that can be captured at the level of individual nodes (dimension *d* = 1) and edges (dimension *d* = 2) have been shown to significantly shape probabilities of fixation (Sood and Redner, 2005; Antal et al., 2006; Broom et al., 2011; Tan and Lü, 2014; Kuo et al., 2021) and are therefore unsuitable for shaping times to fixation while keeping probabilities constant.

**Figure 1:**
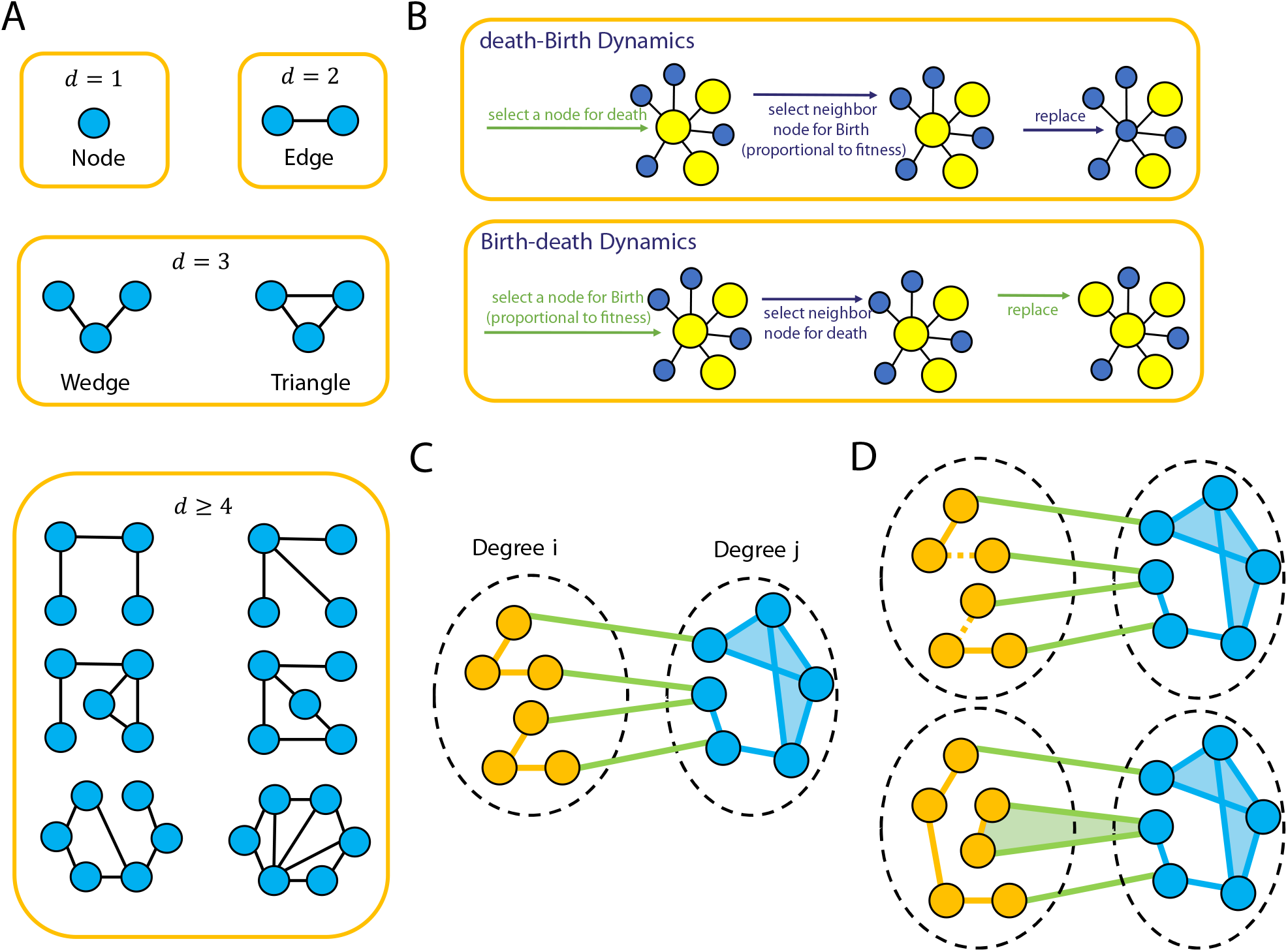
Illustration of the model. **Panel A** illustrates the levels of structural organization in the network. The list of motifs for *d* ≥ 4 is not exhaustive. **Panel B** illustrates the Bd (Birth-death) and the dB (death-Birth) update rules. **Panel C** shows the degree heterogeneous graphs we design, consisting of two groups of nodes with distinct degrees. **Panel D** illustrates the edge swap operation used to tune triangle fractions in the graph, without changing the degree distribution and mixing pattern of the network. Initially, there is no triangle consisting of mixed node degrees. We randomly select two edges of the same type (denoted by color) to be disconnected and nodes that were “parallel” with respect to the two disconnected edges are then connected, thus preserving the number of edges. After the rewiring step, there exists a triangle that connects two yellow nodes and a blue node. The degrees of the nodes and frequencies of edge type, however, are preserved.

Here we systematically explore how higher-dimensional network motifs (*d ≥* 3), and in particular three-dimensional wedges and triangles, shape probabilities and time to fixation of a new mutant in the population and identify network motifs that allow for continuous tuning of times to fixation independent of fixation probabilities. We show that increasing the triangle count of a graph increases times to fixation, without influencing the probability of fixation. We also show that increasing the mean degree of the network not only decreases the time to fixation, but also diminishes the triangle’s ability to shape times to fixation. We find a weak increase with triangle count for the probability of fixation only for highly assortative graphs, but this effect does not undermine the utility of tuning triangles counts for these networks since they are known suppressors of selection (Kuo et al., 2021). In applications where we want to reduce the rate of evolution, increasing time to fixation can achieve the same goal as decreasing the probability of fixation. Collectively, our results suggest that the fraction of triangles is one of the most versatile parameters in controlling fixation acceleration, while maintaining selection amplification.

We apply our algorithms and analytic results to two evolutionary optimization problems and show that the rate at which evolving agents learn to navigate their environment can be tuned near continuously by adjusting the higher-order topology of the agent population. We additionally show that the effects of population structure on the rate of solution discovery are more subtle than previously recognized, and find that decelerators with longer times to fixation are able to reach the optimal solution faster than accelerators in complex solution spaces. Our work highlights that no one population structure is perfect for all optimization tasks, and careful consideration of the problem landscape is necessary for selecting the correct approach. We also show a prevalence of triangles in cellular and social networks, which reinforces the importance of exploring common design principles of real biological networks for engineering highly evolvable artificial systems.

## Model

To study the role of higher-order interactions in shaping evolutionary outcome, we use a Moran birth-death process to track variant frequency changes in a finite population of constant size *N* . We use graphs to represent the population structure. A node in the graph represents an individual in the population, or a homogeneous subgroup of individuals. The network edges are proxies for the local pattern of replacement and substitution: an edge can thus represent spatial proximity or the social architecture of interaction between individuals in the population. The edges could also be interpreted as migration corridors between the homogeneous subpopulations representing the nodes, with the assumption that the timescale of a mutation traveling between nodes is much larger than the time scale of fixation within a node subpopulation. The graphs we consider here are unweighted and undirected.

We compute probabilities and times to fixation of a new mutation *a* that appears in an initially homogeneous population of *A*-type individuals. An individual with the *A* allele is assumed to have fitness one, while an individual with allele *a* has assigned fitness (1 + *s*). We study both Birth-death and death-Birth processes on the networks. These two update rules are equivalent in well-mixed populations, however they have been shown to lead to drastically different long-term evolutionary dynamics (Lieberman et al., 2005; Hindersin and Traulsen, 2015; Kuo et al., 2021). In the Birth-death process, at every time step, a random individual is chosen for reproduction, proportional to fitness. An individual occupying a neighboring node is then randomly chosen for death and replaced by the new offspring. In the death-Birth update, an individual is first randomly chosen for death and is replaced by the offspring of a neighboring node. The neighbors of the vacant node compete for this vacancy and one neighbor is chosen for reproduction with probability proportional to fitness (**Figure 1B**).

Higher-order interactions create graph geometries produced by different patterns of interconnecting nodes. To describe and constrain random graphs by levels of organization in successively finer detail, Mahadevan et al. (2006) previously introduced the mathematical formalism of *dK*-distributions, where *d* denotes the dimension of an interaction involving *d* nodes. Thus, the 0*K*-distribution represents the average node degree of a graph, the 1*K*-distribution is the degree distribution, 2*K* refers to the pattern of interactions, or mixing pattern, between two nodes, 3*K* describes interactions between three nodes, and so on. For 3*K*, there exist two possible connection topologies: 1) wedges, chains of three nodes connected by two edges and their dimension three complement topology, 2) triangles, cliques of three nodes (**Figure 1A**). Here we consider interactions of *d ≥* 3 and particularly focus on the role of 3*K* interactions in shaping probabilities and time to fixation.

To systematically study the role of higher order interactions, the challenge lies in tuning these higher levels of organization while keeping lower levels constant. This is complicated by the fact that, for example, varying the number of triangles in the network also changes degree distribution and network mixing pattern, which we have previously shown to significantly affect probabilities of fixation (Kuo et al., 2021).

We begin by studying the role of higher order interactions in random regular graphs, where all the nodes have the same degree (Steger and Wormald, 1999; Kim and Vu, 2003; Hagberg et al., 2008). This simplifies the problem, since all *k*-regular graphs of fixed degree *k* share the same mixing pattern. We use degree preserving edge swap operations to computationally tune the number of triangles. This allows the tuning of several network properties, while maintaining the degree distribution constant (Taylor, 1981). We build algorithms that combine these approaches with simulated annealing to produce a sampling network generation algorithm. After each edge swap, the fraction of triangles is altered. If the change in the fraction of triangles does not have the correct sign, we accept an edge swap if

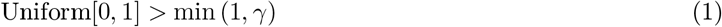

and reject the step otherwise. Here, 1*/γ* is the annealing temperature which controls how stringent the criterion must be and is decreased as the simulation proceeds. Since edge swapping can cause the graph to become disconnected, we use the heuristic outlined in Gkantsidis et al. (2003) to ensure graph connectivity. By saving intermediate graphs, this algorithm generates a set of regular graphs spanning the space of triplet topology. In the **Supplementary Material**, we also present an analysis of the space of *d* = 4 topology.

We expand the analysis to degree-heterogeneous graphs by designing graphs with two distinct degrees (**Figure 1C**), which allows us to seamlessly tune the number of triangles without changing degree distributions and mixing patterns of the graphs. These graphs are designed starting with two random regular networks of size *n*_1_ and *n*_2_ and fixed degrees *k*_1_ and *k*_2_ and we use edge swaps to connect them. Here, the edge swap operation we previously used for the analysis of *k*-regular graphs is unsuitable to tune the fraction of triangles in the network, since that algorithm also alters the graph mixing pattern. Not controlling for the mixing pattern of a graph can potentially lead to erroneous interpretations (see **Supplementary Figure S1**). For studying degree-heterogeneous networks, we use the *dK*-preserving rewiring, which is a generalization of the degree preserving edge swap to higher-order interactions (Mahadevan et al., 2006). **Figure 1D** illustrates a 2*K*-preserving rewiring. We use this variant of edge swapping operation to find graphs with extreme graph properties from all possible graphs of fixed degree distribution and mixing pattern.

For the networks studied, once the network structure is set, we use ensembles of at least 10, 000 Monte Carlo simulations, as well as analytic approaches as described in the next section, to compute the probabilities and times to fixation of the new variant in the population.

## Results

### The role of higher order interactions in networks of uniform degree

We begin by studying the role of higher order interactions in shaping rates of evolution for random regular graphs, where all nodes have the same degree *k*. We describe here the main steps and intuitions behind the analytical approximation for the Birth-death process and provide the complete theoretical analysis in the **Supplementary Material**. We use the diffusion approximation (Kimura, 1962; Crow et al., 1970; Ohtsuki et al., 2006; Sood and Redner, 2005) to estimate the probability and time to fixation of the new *a* mutant as it appears on a random node of the network. We denote by *p_a_* the mutant frequency in the population. There are three possible types of edges in the network: *AA*, *Aa*, *aa* and we denote their frequencies as *p_AA_*, *p_Aa_*, *p_aa_*. These frequencies can be expressed in terms of *p_a_* and *p_Aa_*, with *p_AA_* = (1 *− p_a_*) *− p_Aa_* and *p_aa_* = *p_a_ − p_Aa_*. At every time point, *p_a_* either remains the same, increases by 1*/N*, or decreases by 1*/N* . We can write the mean and variance of the change in node frequency as

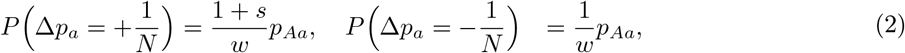

where *w* is the mean fitness of the individuals in the population.

Since mutant frequency only changes when a replacement event occurs between edges connecting a wildtype and mutant, the whole dynamical system can be described by changes in *p_a_* and *p_Aa_*.

The expected change in the frequency of the mutant, *p_a_*, is 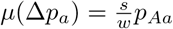. Using pair approximations, we compute the expected change in *Aa* edge-type frequency, *p_Aa_*:

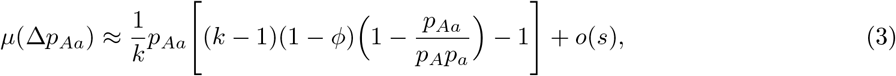

where *ϕ* is the transitivity or the number of closed triangles divided by the total number of triples.

Observe that the expected change in mutant node frequency is on the order of the selection coefficient *s*, while the expected change in *Aa* edge frequency is independent of *s*. Therefore, in the limit of diminishing selection, the dynamics of the edges are faster than those of the nodes and we can assume that the edge frequencies are at equilibrium in the timescale of the node dynamics. We can write *p_Aa_* at equilibrium

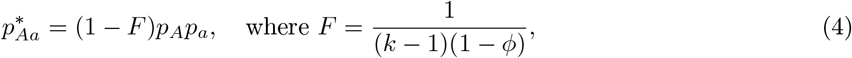

and using this approximation of *p_Aa_*, the Kolmogorov equation reduces to depend on *p_a_* alone. The probability of fixation of the *a* allele for any initial mutant frequency *P* (*p_a_*) is approximated by the solution to the equation:

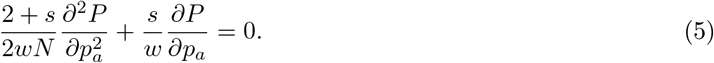

This equation is identical to that corresponding to a well-mixed population, as expected for *k*-regular graphs (Lieberman et al., 2005). The probability of fixation of the new mutant is therefore independent of the higher-order configuration of the nodes of the network.

While higher-order motifs may not shape probabilities of fixation, they can drastically change the time to fixation of the new variant. The time to fixation, conditional on fixation, *T* (*p_a_*) can be found by solving

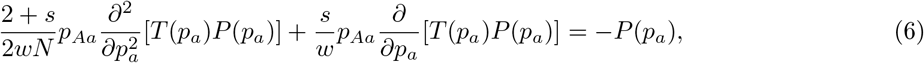

and can be written as

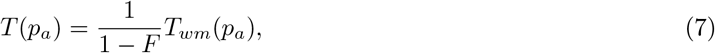

where *T_wm_*(*p_a_*) is the conditional fixation time for the well-mixed model, approximated by (2+*s*)*s^−^*^1^*N* log *N*, for *s >* 0 (Antal and Scheuring, 2006).

The analytic approximation provides intuition into the dynamics of the model. The equilibrium frequency of *Aa* edges is minimized in a population of all *A* or all *a* individuals (**Figure 2A**) and as the frequency of *a* increases, the frequency of *Aa* edges quickly converges to a constant, (1 *− F*), times the product of the frequencies of *A* and *a*. This constant decreases with the fraction of triangles *ϕ* (**Figure 2B**) and this decrease leads to higher times to fixation for the new variants in the population. Parameter *F* in equation (7) is analogous to measures of inbreeding due to population structure. Intuitively, rewiring a network of fixed degree distribution to introduce triangles (**Figure 2C**) increases the probability that two nodes with the common ancestor are connected, thereby reducing the number of edges connecting nodes of different genotypes.

**Figure 2:**
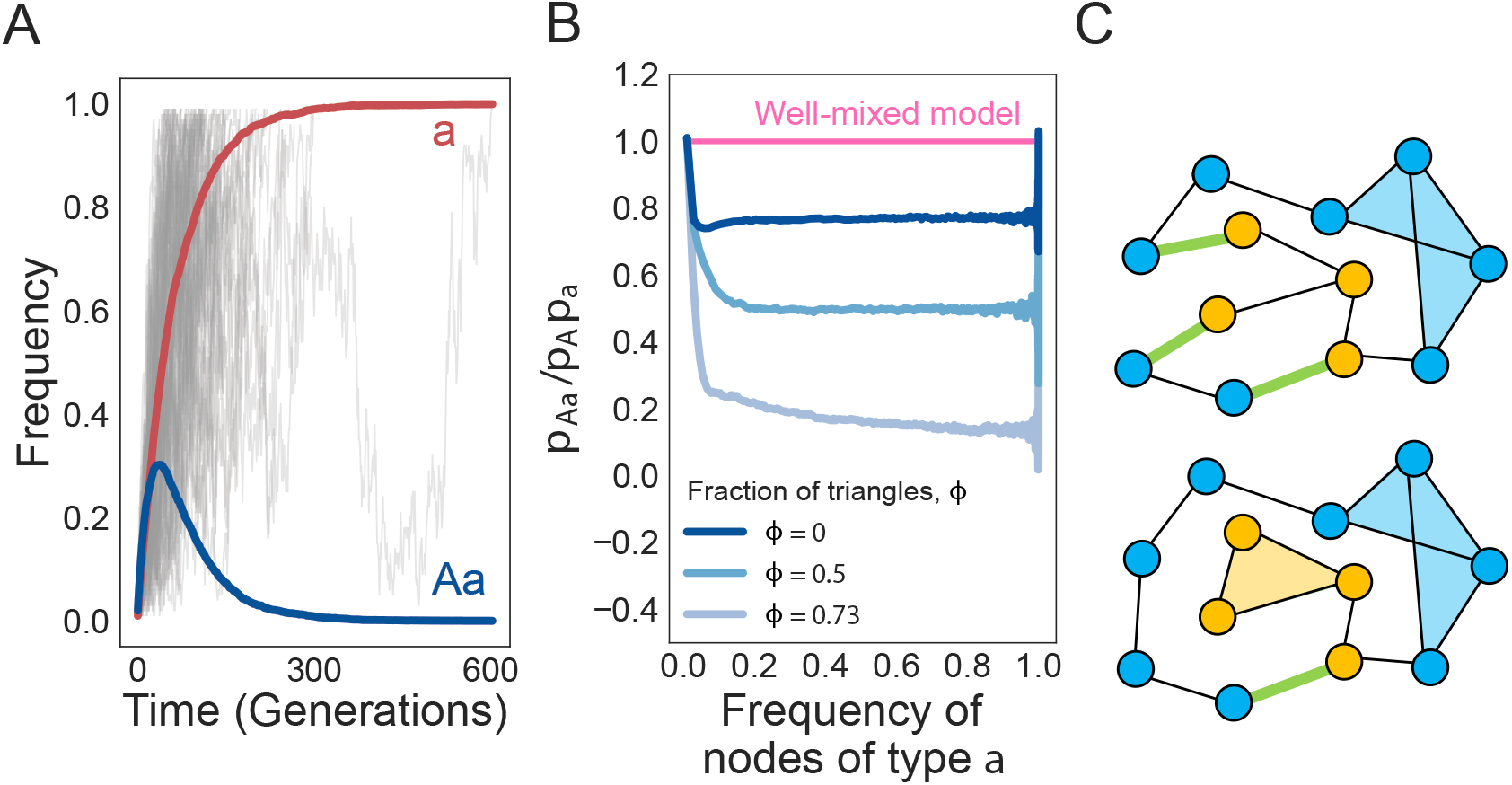
Triangles reduce the equilibrium number of *Aa* edges in the graph. **Panel A** shows the time trajectory of mean frequencies of *a* nodes and *Aa* edges from ensembles of Monte Carlo simulations (individual trajectories shown in gray) on a 5-regular graph of size 100 with 0 triangles. Only trajectories that fix in *a* were used. **Panel B** shows the ratio between frequency of *Aa* and the frequency of *a* nodes on graphs with various fractions of triangles *ϕ*. All results were obtained from 1000 replicate simulations. **Panel C** shows two graphs with the same degree distribution. The top graph has two triangles while the bottom one has three triangles. The number of open edges in green (edges that connect pairs of different genotype) is reduced when triangles are introduced.

We show the accuracy of the analytic approximation for random regular networks in **Figure 3**. The fixation probability is independent of the node degree (**Figure 3A**) and, as the fraction of triangles *ϕ* increases, time to fixation increases as *F* approaches 1 (**Figure 3B**). Analogous to the amplification parameter in quantifying the probabilities of fixation (Kuo et al., 2021), we define the acceleration factor as the time to fixation in a well-mixed population over the time to fixation in a structured population. For *k*-regular graphs, the acceleration parameter is approximately (1 *− F*), which increases as the mean degree increases and the fraction of the triangles decreases (**Figure 3C**). Our analysis suggests that *k*-regular graphs are not accelerators of evolution, since the acceleration parameter cannot exceed one. We also show that our approximation of time to fixation holds for a wide range of selection strengths (**Figure 3D**), even though in our derivation we assume weak selection.

**Figure 3:**
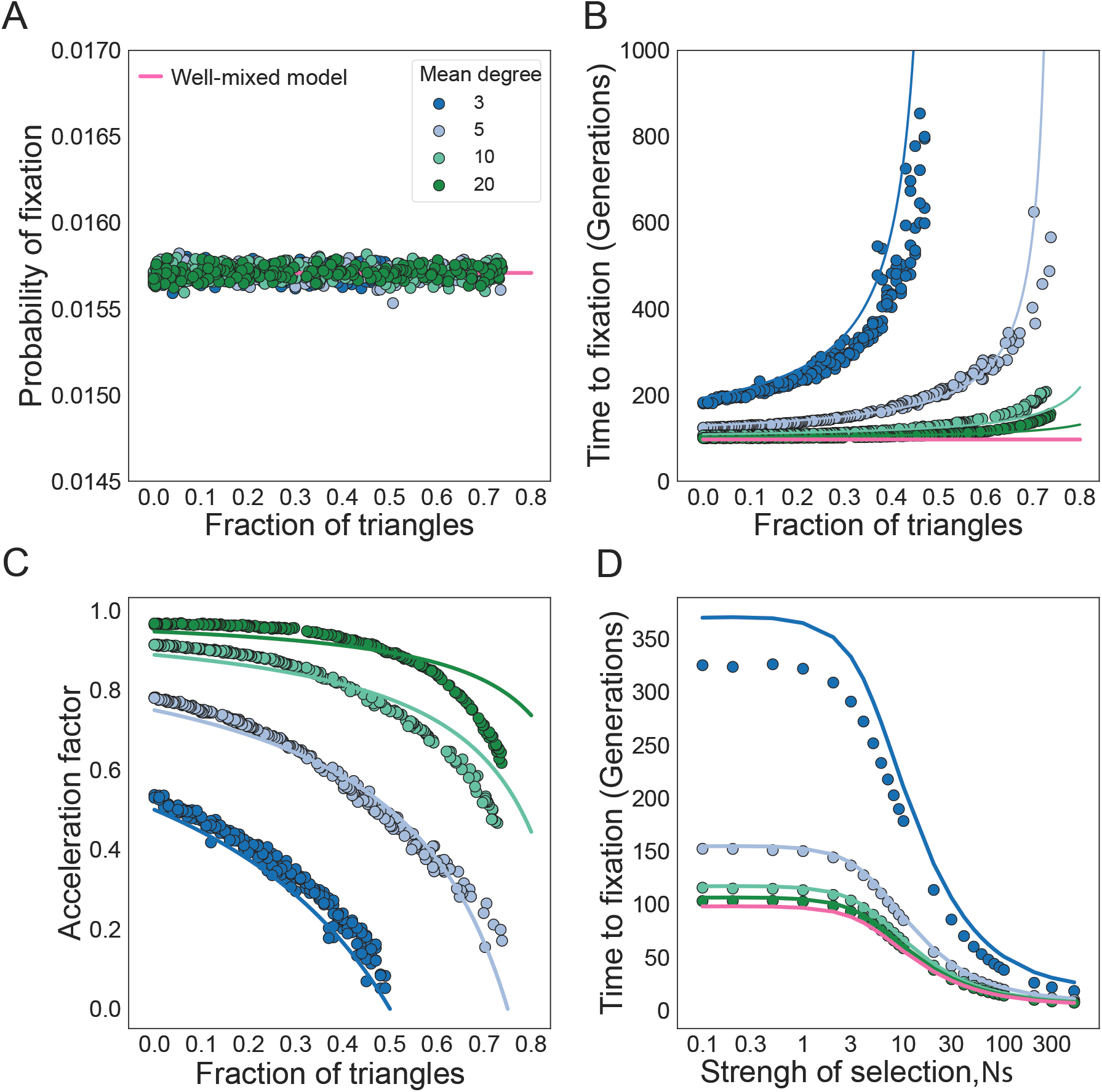
The fraction of triangles increases time to fixation in degree uniform graphs. The dots represent ensemble averages across 10^6^ replicate Monte Carlo simulations, while the lines represent our analytical approximations. Here the degree distribution is held constant as we vary the fraction of triangles in the graphs. *N* = 100 and *s* = 0.01, such that *Ns* = 1. The color indicates the mean degree of the network, as in the legend. The fraction of triangles in the graph is tuned using edge swapping operations. **Panel A** shows the fixation probability, **Panel B** shows the fixation time, **Panel C** shows the acceleration factor as a function of the fraction of triangles, and **Panel D** shows the time to fixation as a function of selection strength.

### Higher-order interactions increase time to fixation in degree-heterogenous networks

For degree-heterogeneous networks where the node assortativity is zero, the fraction of triangles similarly increases the time to fixation (**Figures 4B** and **4D**), while having a negligible effect on the probability of fixation (**Figures 4A** and **4C**). We vary mean and standard deviation in degree independently and observe that the effects of triangles are more pronounced in networks of small mean degree and high standard deviation in degree.

**Figure 4:**
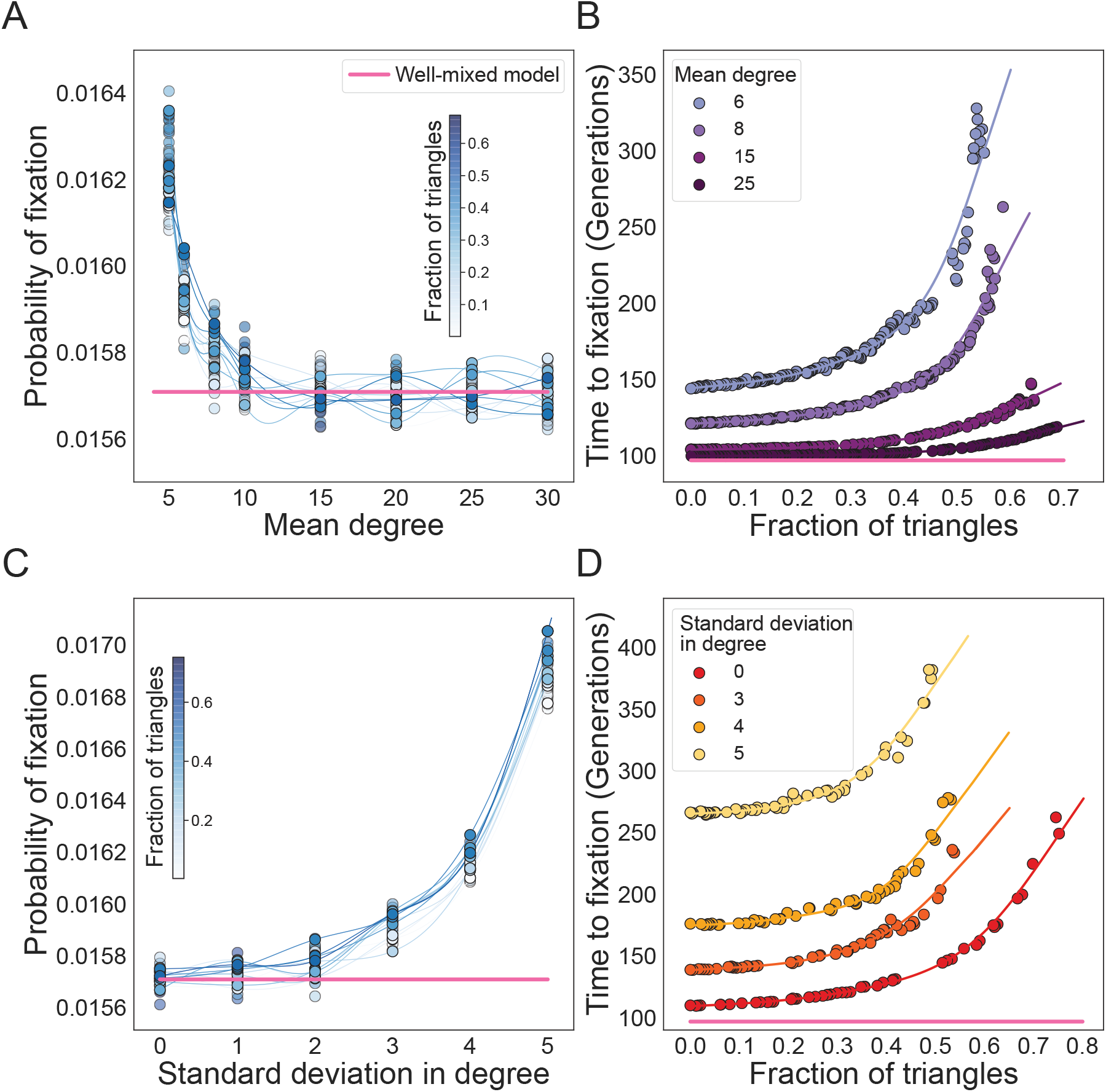
Higher-order interactions increase time to fixation in degree heterogenous graphs with zero assortativity. Here the graph mixing pattern is constant, with assortativity equal to zero, *N* = 100 and *s* = 0.01, such that *Ns* = 1. **Panel A** shows the fixation probability as a function of the fraction of triangles and mean degree of the graph. **Panel B** shows the time to fixation, and the colors indicate the mean degree of the network as in the legend. Standard deviation in degree equal to two. **Panel C** shows the fixation probability as a function of the fraction of triangles and the heterogeneity in the degree of the graph. **Panel D** shows the time to fixation. For **Panels C** and **D**, mean degree equal to eight. The dots represent ensemble averages across 10^6^ replicate Monte Carlo simulations, while the lines represent cubic spline regression.

As we increase network assortativity, the frequency of higher-order interactions changes both the probability and the time to fixation of the new allele *a*. To assess the role of assortativity, we design graphs with fixed degree distribution and vary the mixing pattern using the algorithm from Kuo et al. (2021). **Figure 5A**, shows that the fraction of triangles has negligible influence on fixation probability for disassortative and weakly assortative graphs. However, as we increase assortativity, the probability of fixation increases for networks with many triangles. This observation can be explained by considering the expected number of *Aa* edges in the network. Similar to the case of *k*-regular networks, triangles reduce the number of *Aa* edges in the network (**Figure S2**). The key difference here is that regular graphs only have one type of *Aa* edges, while more complex networks have multiple types of *Aa* edges as defined by the various degrees of the nodes they connect, *A_i_a_j_*. If the *A_i_a_j_* edges are reduced by the fraction of network triangles by roughly the same amount, their effects cancel out and the probability of fixation remains unchanged. In the case of highly assortative graphs however, *A_i_a_j_* edge frequencies tend to decrease more for edges that connect nodes of low degree. The same behavior is observed across a wide range of graph families (see **Supplementary Figures 5**, **6**, and **7**). The increase in the time to fixation due to graph triangles is amplified by network assortativity (**Figure 5B**). Intuitively, this is because the number of *A_i_a_j_* edges is reduced even more in assortative graphs. In **Figure 5C**, we show that the deceleration in fixation for networks with a fixed fraction of triangles is constant across biologically relevant selection ranges *Ns* from 0.1 to 100. This deceleration is maintained across a wide range of selection, suggesting that these networks are not just piece-wise decelerators.

**Figure 5:**
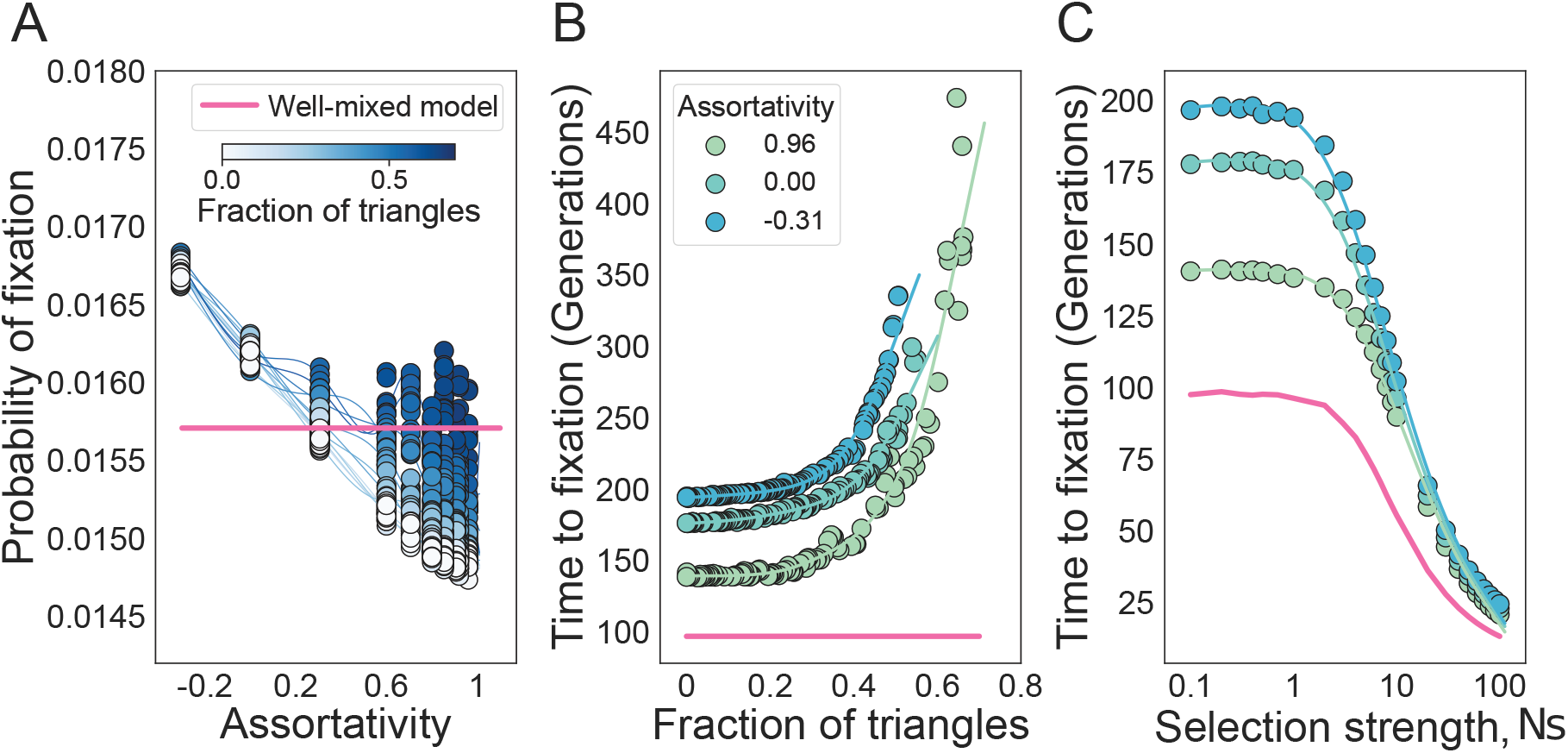
The probability and time to fixation in graphs with strictly positive assortativity. Here the degree distribution of the network and its assortativity is held constant, as we vary the fraction of triangles in the graphs. *N* = 100 and *s* = 0.01. The degree distribution is 50% nodes of degree 12 and 50% nodes of degree 4, such that the mean is 8 and the standard deviation is 4. **Panel A** shows the probability of fixation as a fraction of triangles and assortativity of the graph. **Panel B** shows the time to fixation and the colors indicate the assortativity of the network, as in the legend. **Panel C** shows how the effect of the strength of selection for graphs with different assortativity and a fixed fraction of triangles equal to zero. The dots represent ensemble averages across 10^6^ replicate Monte Carlo simulations, while the lines represent cubic spline regression.

**Figure 6:**
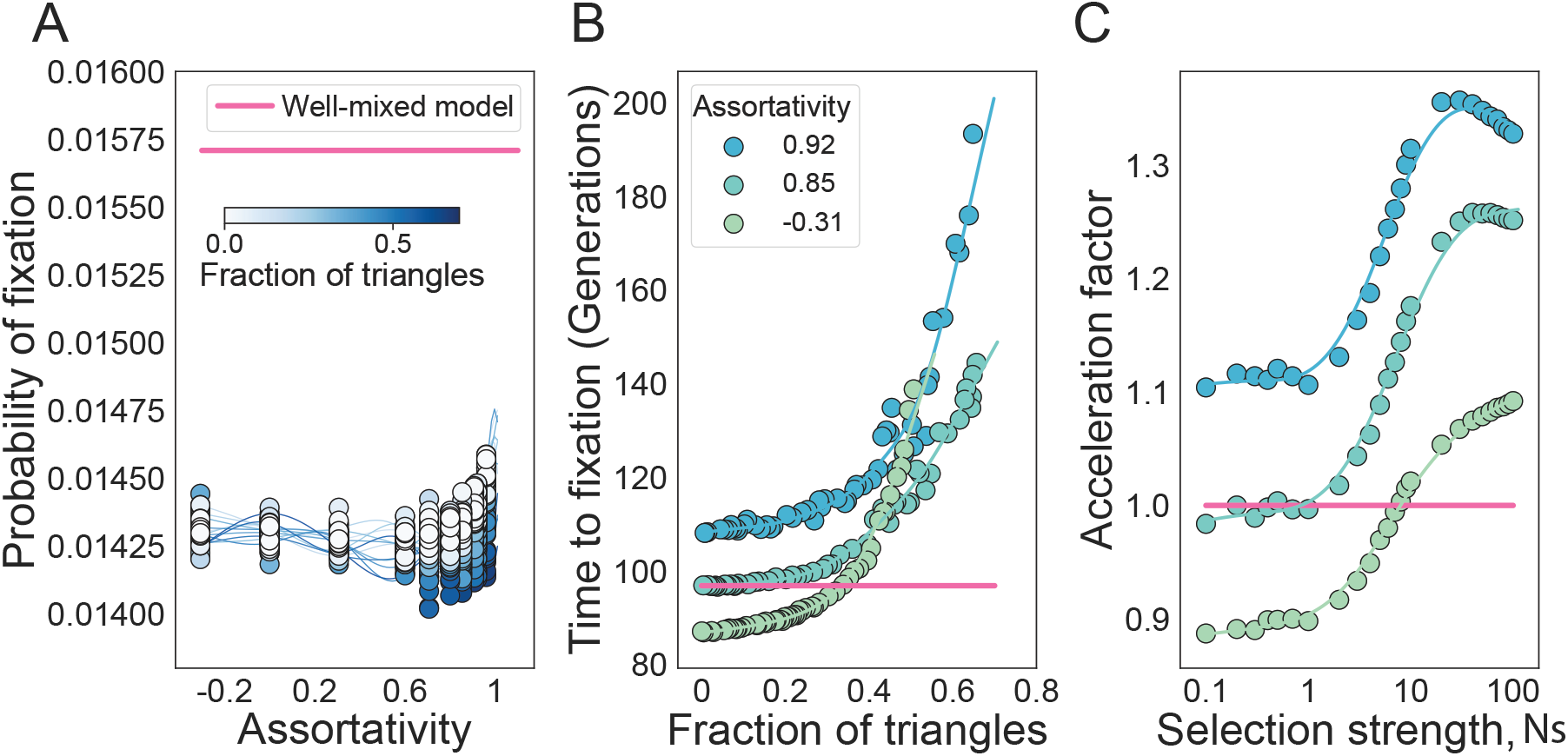
Population structures that accelerate the time to fixation are abundant, but only under death-Birth dynamics. **Panel A** shows the probability of fixation with respect to assortativity and fraction of triangles of the graph. **Panel B** shows the time to fixation and the colors indicate the assortativity of the network, as in the legend. **Panel C** shows the effect of the strength of selection for graphs with different assortativity and a fixed fraction of triangles equal to zero. The dots represent ensemble averages across 10^6^ replicate Monte Carlo simulations, while the lines represent cubic spline regression. Here the degree distribution of the network and its assortativity is held constant, as we change the fraction of triangles in the graphs. *N* = 100 and *s* = 0.01. The degree distribution is 50% nodes of degree 12 and 50% nodes of degree 4, such that the mean is 8 and the standard deviation is 4.

The role of triangles is invariant to the update process: under a death-Birth process, the fraction of triangles has minimal effects on the probability of fixation (**Figure 6A**) and the time to fixation similarly increases with an increased fraction of triangles (**Figure 6B**). However, we find that graphs with low assortativity and a low fraction of triangles exhibit a faster time to fixation compared to the well-mixed model. This acceleration, however, is not stable with respect to selection. **Figure 6C** shows that the acceleration factor increases as the strength of selection increases, turning accelerators into decelerators. While all graphs under the Birth-death process are decelerators of evolution, we find that a piece-wise accelerator of fixation is prevalent under the death-Birth update rule (**Figure 6**).

### Optimizing network structure in evolutionary algorithms applications

Our results suggest that network higher-order topologies could have important applications in evolutionary optimization problems. We first explore the role of network 3D motifs when evolving an agent to optimize reward in the cart-pole balancing problem (Anderson, 1987). We use a variation of the evolutionary algorithm on graphs described in Bryden et al. (2006). As illustrated in **Figure 7A**, individuals in the population are represented by neural network weights. A neural network receives the state of the environment as input and outputs an action. If the neural network performs the right action, then that individual gets a positive reward from the environment. The fitness of the individual is calculated by its rank in the population, where the best performing agent is assigned fitness of (1 + *s*), decreasing linearly to (1 *−s*). Such a selection scheme is called ranked selection which maintains constant selection pressure on the population, independent of the problem (Baker, 1985). The population evolves under the Birth-death process. We use 5-regular graphs of size *N* = 100 and a selection strength of *s* = 1 such that *Ns* = 100. Every time a node reproduces, either the parent or the offspring node mutates by adding a Gaussian noise vector to the neural network.

**Figure 7:**
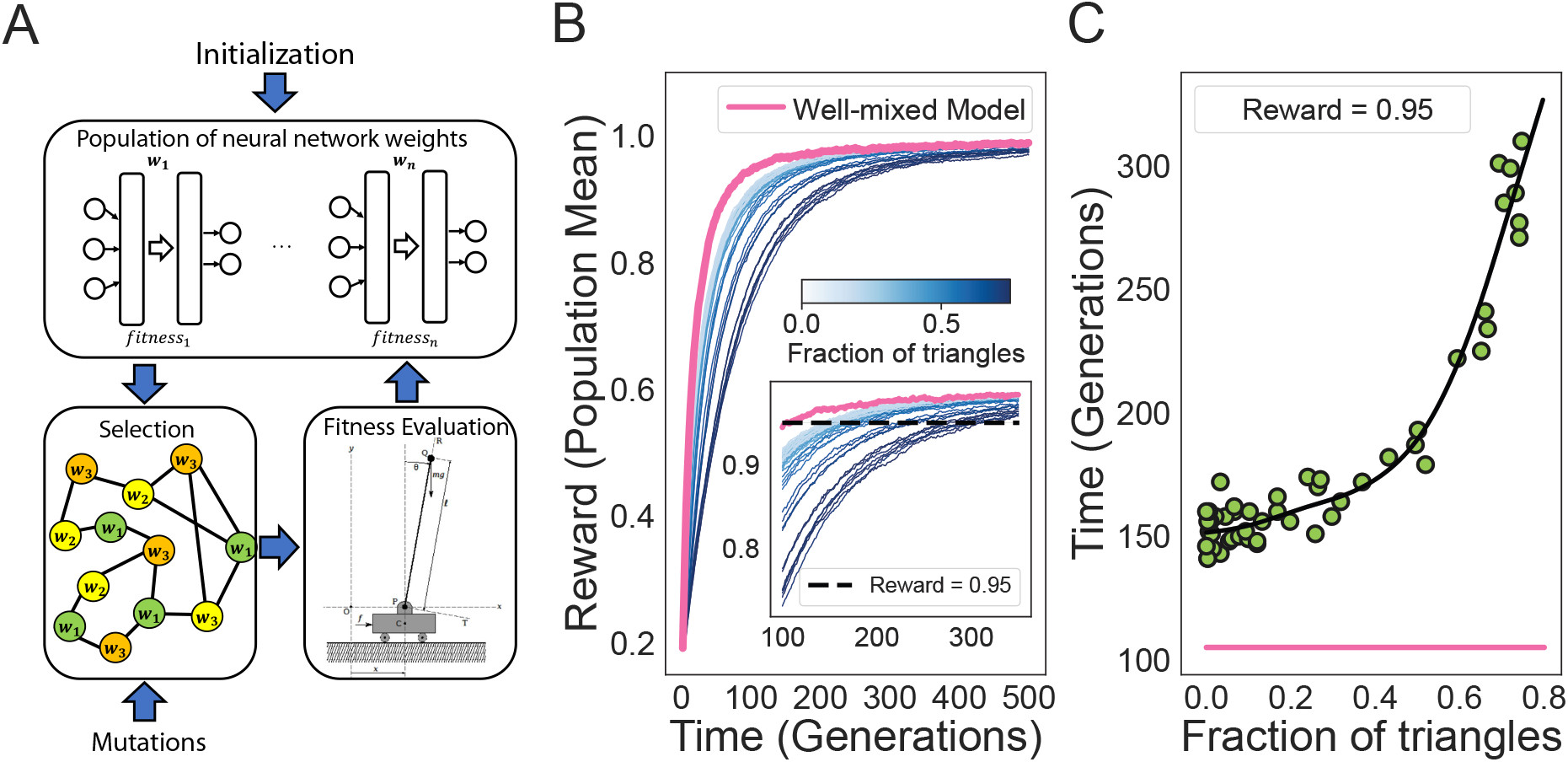
Effects of network topology on rates of convergence in the cart-pole balancing problem. The population structure used in the simulation have mean degree of five and varying transitivity *ϕ*. The results are averaged from 500 replicate simulations. **Panel A** shows the illustration of the evolutionary algorithm using structured populations. In **Panel B** the lines represent average trajectories of the reward received in the Cart-pole problem. The outer plot shows the average reward received by the population, while the inner plot shows the time it takes for each population to reach a high reward of 0.95. In **Panel C** the dots represent the average time to reach a reward of 0.95 as a function of the higher-dimensional population structure.

Increasing the fraction of triangles in the network reduces the rate that the population discovers the optimal agent (**Figure 7B**). The population structure with the highest number of triangles has the lowest rate of convergence to an optimal reward of one, and the rate of convergence increases as the fraction of triangles decreases. **Figure 7C** shows the time it takes for each topology to reach a normalized reward of 0.95. As expected, this time increases as a function of the fraction of triangles in the network.

While triangles in a network may be detrimental in optimization problems on smooth fitness landscapes, in problems where the optimization landscape is rugged, increasing the fraction of triangles leads to a population with an increased rate of solution discovery. The second application we consider is finding the global minimum of the Rastrigin function (Rastrigin, 1974). We invert the function, turning the problem into a maximization problem. The Rastrigin function has tuneable ruggedness, which makes the gradient-based optimization used in convex optimization unsuitable (**Figure 8A**). Similar to our first application, each individual in the population is represented by a vector in the domain of the Rastrigin function and fitness is mapped to the population ranking of the function value corresponding to the individual. The population reproduces on networks under the Birth-death process and Gaussian mutations occur randomly to change the offspring or the parent node. The population structure with the lowest number of triangles has a higher rate of initial convergence (**Figure 8**B), but it is soon overtaken by populations with higher numbers of triangles in the network (**Figures 8C** and **D**).

**Figure 8:**
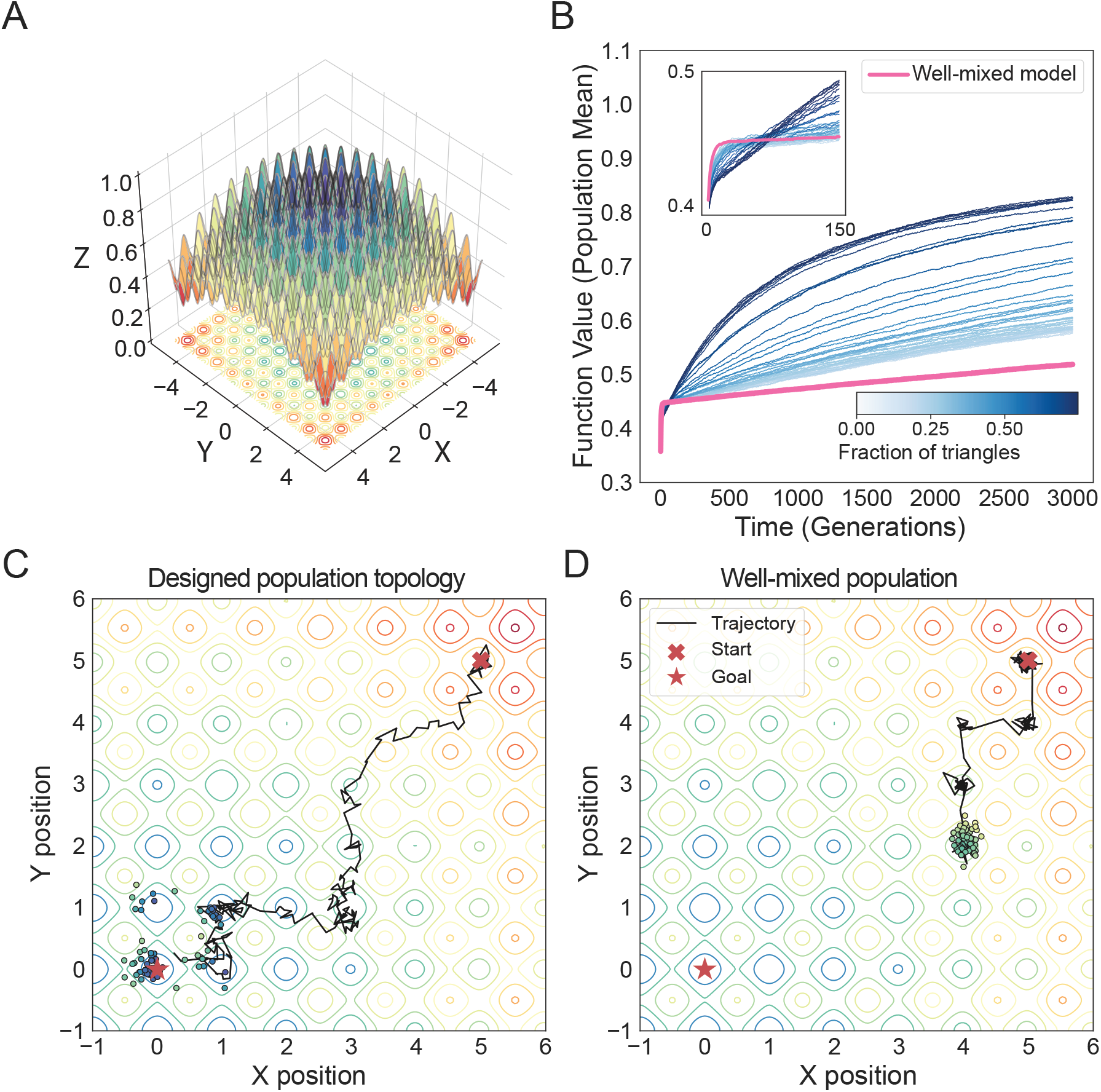
Advantage of increased fixation time in complex optimization problems. The population structures used in the simulation have mean degree of five and varying transitivity *ϕ*. The results are averaged from 500 replicate simulations. **Panel A** illustrates the Rastrigin function, used as test function. **Panel B** shows mean solution trajectories. **Panel C** shows the population trajectory and the distribution of solutions in function space for the network population, while **Panel D** shows the population trajectory the distribution of solutions in function space for a well-mixed population.

## Discussion

Modern approaches to training agents to reach human-level performance utilize Deep Neural Networks, which are typically optimized using gradient-based methods, such a back-propagation. Recent successes in deep reinforcement learning that train agents end-to-end from high dimensional data sparked further development of evolutionary algorithms and recent work suggests a simple Genetic Algorithm (GA) is possible to optimize DNNs and that GAs are a competitive alternative to gradient-based methods when applied to Reinforcement Learning tasks such as learning to play Atari games, simulated humanoid locomotion, or deceptive maze problems (Such et al., 2017). In fact, evolution and gradient-based methods are not competing approaches to optimizations and a hybrid approach will most likely become the best implementation (Stanley et al., 2019).

To design artificial populations with accelerated (or decelerated) rates of evolution for applications in directed evolution and evolutionary optimization, we must first build a systematic understanding of how the spatial arrangement of a population shapes evolutionary dynamics. Here we study how higher-order network topologies - and in particular 3-dimensional structural motifs - shape the probabilities and times to fixation of new variants in a population. Our results show that triangle count is the most effective network property for tuning temporal dynamics. Our results inform on the design of population structures that allow us to control evolutionary spread of optimal solutions and can be applied to a wide range of optimization problems. We show that fast fixation time is not always ideal and can lead to slow solution discovery when the optimization landscape is rugged.

Initial spatial evolutionary algorithms on lower-dimensional exhibited a longer polynomial take-over time compared to the logistic growth rates in algorithms based on well-mixed populations (Giacobini et al., 2005). In those studies, the takeover time, equivalent of our time to fixation here, is controlled through the size and shape of the neighborhood in the lattice, as well as the population update rules. Empirical studies on highly symmetric or regular population structures suggest that the optimal graph diameter increases and the optimal average degree decreases with the complexity and difficulty of the fitness landscape (Bryden et al., 2006). Adding the use of the average number of contacts in the population offers even more flexibility over update rules in controlling the dynamics of mutational spread. However, the average number of contacts takes a discrete integer value in regular graphs, and our analysis suggests the existence of large gaps in the spectrum of fixation time, especially when mean degree is low. This leads to limited ability to tune desired evolutionary rates, in particular in cases where slowed spreading is required. Analysis of scale-free interaction networks suggests takeover time can also be controlled by fine-tuning the scaling exponent *γ* or network assortativity *r* (Payne and Eppstein, 2009). While these quantities are continuous, they are known to also significantly affect the probability of fixation as well, leading to trade-offs and complex behavior in the effective rate of evolution. The fraction of triangles of a network allows finer evolutionary tuning because it is a nearly continuous variable, in the limit of large population size and it keeps probabilities of fixation constant.

Our theoretical treatment is limited to studying probabilities and times to fixation, but more factors contribute to the evolution of these systems. For example, evolutionary algorithms purposely select elevated mutation rates to have multiple mutants simultaneously in a population. This allows the algorithms to search different solutions simultaneously. Different clones compete and can interfere with the fixation of one another (Gerrish and Lenski, 1998). Effective traversal of the fitness landscape also relies on the existence and maintenance of mutation islands. Fixation time might not be the most direct quantity that describes the clonal dynamics in such a population and tools that study clonal interference could potentially also provide additional insights into the problem. Furthermore, future studies that modify four or higher node structures independently could also reveal their underlying roles in evolutionary dynamics.

We further apply our theoretical results to understand the topological organization of biological and social networks of organization. We study the hematopoietic stem cell organization in the bone marrow as a graph where each node is a niche and edges connect niches within biological ranges of interaction (Kuo et al., 2021). We find that the cellular spatial organization in the bone marrow maximizes the number of triangles in the network (**Figure S8A**). We plot triangle count against the average triplet count of the networks. The true inferred network is shown in blue, on a background of a range of network topologies, as shown in gray. The true networks have higher triangle counts than the tuned graphs, even when the tuning algorithm is designed to maximize the fraction of triangles. Previous work also suggests that spatial networks tend to have large numbers of triangles, since connection costs favor the formation of cliques between spatially close nodes and thus increase the network clustering coefficient (Barthélemy, 2011). One example is the brain anatomical and functional networks which exhibit dense local clustering or cliquishness of connections between neighboring nodes. This minimizes wiring costs while supporting high dynamical complexity. Loss of cliquishness is a signature of diseases that impairs brain function such as Alzheimer (Supekar et al., 2008) and schizophrenia (Liu et al., 2008). The structure of the tissue population also provides insights into how normal cells traverse the mutational landscape towards cancer. The evolutionary landscape is complex, likely comprising of both smooth uphill paths, as well as rugged detours to fitness peaks corresponding to initial neoplasm. A higher number of triangles hinders the traversal of smooth fitness landscapes, while accelerating passage through rugged terrains.

We can also study triangle topologies in culturally evolving populations and use Facebook friendship networks across 20 universities (Rossi and Ahmed, 2015). Since most of the social networks in the dataset are disconnected and exhibit community structure, we extract the connected communities from the network using a community structure detection algorithm and keep the largest community (Newman, 2003). Triad closure, where there is a heightened probability of two people becoming friends if they have one or more other friends in common, promotes the formation of triangles in social networks (Rapoport, 1953) and has been empirical observed in evolving social networks (Newman, 2001). We find that social networks also maximize the number of triangles in the network (**Figure S8B**).

There exist therefore strong design similarities between social and biological networks and this suggests there are more design principles in real-world networks to be explored for engineering of optimal artificial populations.

## Acknowledgments

We gratefully acknowledge support from the United States-Israel Binational Science Foundation (award no. 2019266) and from the NIH T32 training grant (no. T32 EB009403). This research was done using resources provided by the Open Science Grid, which is supported by the National Science Foundation award 1148698, and the U.S. Department of Energy’s Office of Science.

## Supplementary Material

### 1 Analytic approximation for *k*-regular graphs

Here we present a full description of our analytic approach for *k*-regular graphs under the Birth-death update rule. A population of size *N* of *A* individuals is distributed over the nodes of a *k*-regular graph (each node has *k* neighbors). Using the diffusion approximation, we compute the probability and time to fixation, conditional on fixation, of a new mutant *a* appearing in a random node on the graph. We adopt an approach similar to Ohtsuki et al. (2006) and use pair approximations to write higher-order interaction dynamics as approximations of lower-order ones.

At every time step, let *p_a_* and *p_A_* denote the frequencies of the mutant *a* and wild-type *A* in the population. Let *p_aa_*, *p_aA_*, *p_Aa_* and *p_AA_* denote the frequencies of the edge types and let *p_Y |X_* denote the probability of observing an *XY* edge given the first node is type *X*. Here, *X* and *Y* can be *A* or *a*. Note that *p_Aa_* = *p_aA_*, *p_aA_* = *p_a|A_p_A_* and *p_A|X_* + *p_a|X_* = 1.

Let 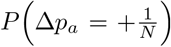 and 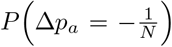 denote the probabilities that an individual changes its allelic type towards or away from the mutant type *a*. Since mutant frequency only changes when a replacement event occurs between edges connecting a wild-type and mutant, the whole dynamical system can be described by changes in *p_a_* and *p_Aa_*.

The mutant frequency increases by 1*/N* in the population when a mutant *a* is selected for reproduction and a neighbor *A* is selected to be replaced. Therefore, *p_a_* increases by 1*/N* with probability

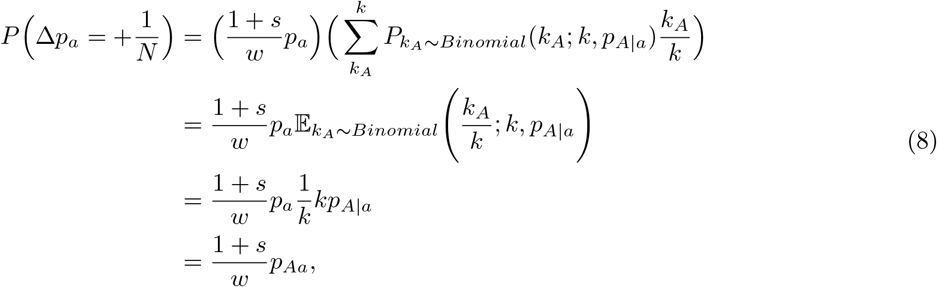

where *k_A_* and *k_a_* denote the number of *A* and *a* neighbors of the focal mutant node and *w* represents the mean fitness of the population. We have *k_A_* + *k_a_* = *k*. The term (1 + *s*)*p_a_/w* corresponds to the probability of selecting a mutant *a* for reproduction. The rest of the term corresponds to the probability of selecting a wild-type *A* to be replaced.

Similarly, we can write the probability that the mutant frequency decreases by 1/*N*,

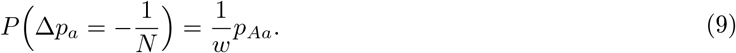

Therefore, the first and second moment of the change in frequency of the mutant allele *a*, at every time step, can be written as:

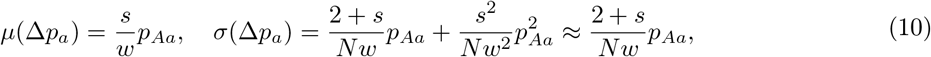

in the limit of weak selection.

We next compute the first and second moment of the change in frequency of the edges of type *Aa*, *p_Aa_*. The number of *Aa* edges only changes when the mutant *a* replaces a wild-type *A* and vice versa.

Firstly, when *a* replaces an *A*, the number of *Aa* edges decreases by one, but that is not the only edge type-changing. We also need to consider edges connecting the death node *A* to nodes other than the birth node *a*. Let us denote these nodes *a*’ or *A*’ . The *AA*’ edges (of number *k_A_*) connecting the node to be replaced *A* to neighboring *A*’ nodes become *aA*’ edges and all *Aa*’ edges connecting neighboring *a*’ nodes to the death node *A* become *aa*’ edges. In total, we lose (*k_a_ −* 1) *Aa* edges and gain *k_A_ aA* edges. The probability of this event is the product of the probability of an *a* node replacing an *A* node and the probability of the dead *A* node having exactly *k_a_ a*’ neighbors and *k_A_ A*’ neighbors. The probability that the frequency of *Aa* edges increases by 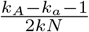 can be written as:

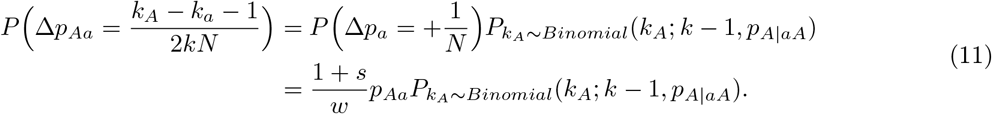

Similarly, we can write the equivalent probability, for the event of a wild-type node *A* replacing a mutant node *a*:

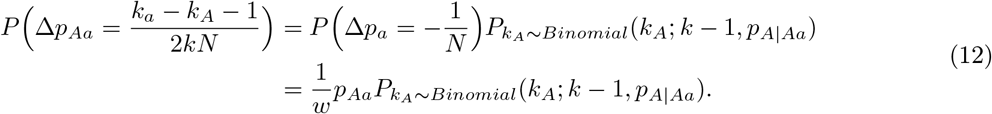

The expected change in frequency of the *Aa*-type edges is sum of the expected change under these two possible replacement events. The expected change under the first event (*a* replaces *A*) is

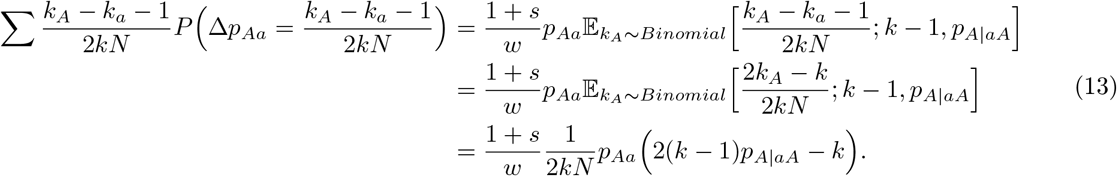

Note that the probability of observing a particular configuration of the death node’s neighbors is conditioned on observing the *Aa* edge between the birth and death nodes. Hence, *k_A_* + *k_a_* = *k −* 1 here.

The expected change under the second event (*A* replaces *a*) is

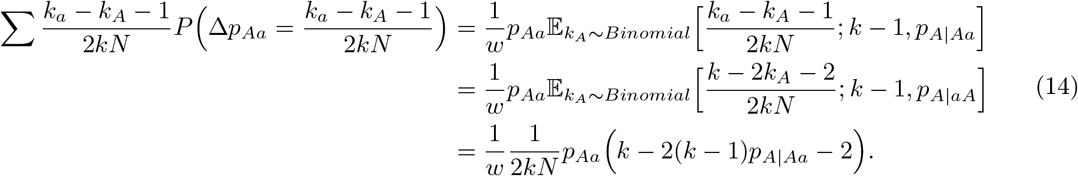

In the limit of diminishing selection we keep only terms that are independent of *s*. The expected change in frequency of the *Aa*-type edges is

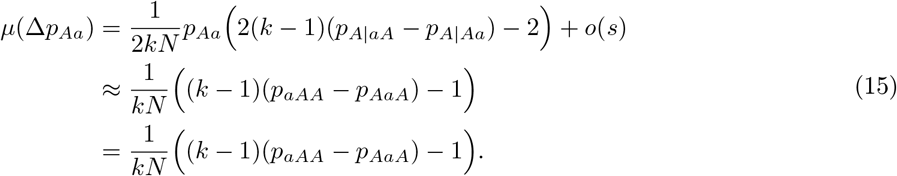

From equation (15), we observe that the expected change in the frequency of *p_Aa_* depends on the frequency of higher-order interactions, specifically the triples *p_AAa_*, *p_AaA_*. Similarly, the change in frequency over time of the triples depends on frequency of quartets. By induction, the change in frequency of (*n−* 1)-tet depends on changes in frequencies of (*n*)-tet until *n* is equal to the population size *N* . This chain of dependencies can be simplified by the use of pair approximations, writing frequencies of (n)-tetas functions of frequencies of the lower levels of network organization.

Using approximations from (House and Keeling, 2011), we write the frequencies of arbitrary triples of nodes *XY Z* as

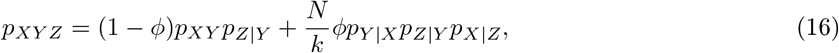

We can rewrite equation (15) as

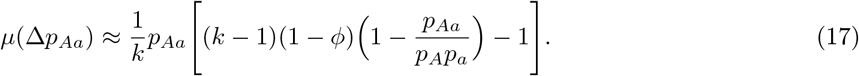

We therefore need to keep track of frequency changes of the nodes and edges only.

We further observe that the expected change in node frequencies is on the order of *s* and the expected change in edge frequencies is independent of *s*. In the limit of diminishing selection, the dynamics of the edges are faster than that of the nodes and we can assume that the edge frequencies are at equilibrium in the timescale of the node dynamics. We write the equilibrium frequency of *Aa* edges as

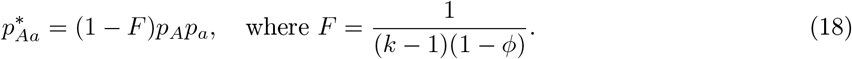

p^∗^_Aa_ = 0 is also an equilibrium frequency, but this state is only reachable for non-zero *p_a_* in disconnected graphs. Here *F* is controlled by both the mean degree and the fraction of triangles of the network. Increasing the mean degree or decreasing the triangle count decreases *F* and increases the equilibrium number of *Aa* edges in the network.

We can now write the Kolmogorov backward equation for the combined node and edge dynamics and equation (18):

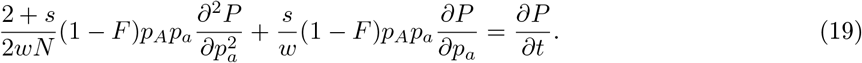

The probability of fixation given any initial mutant frequency is found by solving the KBE for zero

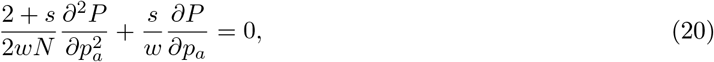

which is identical to the KBE for the well-mixed model, meaning the probability of fixation is unaltered by the topology in regular graph.

The time to fixation conditioned on fixation of the mutant, on the other hand is, is not invariant to graph topology. We calculate the conditional fixation time from results on exit time in stochastic systems (Gardiner, 2009). The time to fixation is given by solving the following equation for *T* :

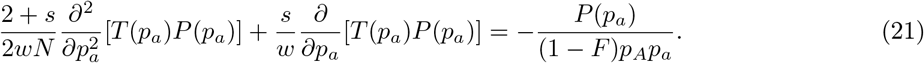

The solution of this equation is

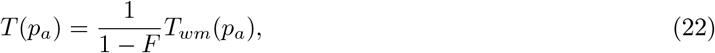

Therefore, the time to fixation in regular graphs is a constant times the fixation time in the well-mixed model, under weak selection. Our results show that increasing the triangle count increases the constant *F* and decreases the equilibrium number of *Aa* edges in the network, which translates into increased fixation time for the new mutant *a*.

### 2 Effects of 4-node Motifs on Evolutionary Dynamics

We also explore the effects of 4-node motifs on the evolutionary dynamics of the new mutation *a*. The dK-preserving edge-swap algorithm is only defined for *d* = 3. There is no edge-swap operation that is guaranteed to preserve the triplet distribution and the fraction of triangles in the network. However, in degree uniform graphs, edge swap does not affect the triplet distribution since all nodes have the same degree; only the ratio of open triples and triangles is altered. We tune the fraction of 4-node motifs (**Supplementary Figure S3 A**), by randomly selecting edges for a 3K edge-swap and only accepting the change if the resulting graph contains the same number of triangles. We combine this with an optimization algorithm to get a range of graphs with different motif counts. The resulting algorithm can not, however, tune the occurrence of 4-motifs independent of other 4-motifs. The relationships between 4-cliques and other 4-motifs are shown in **Supplementary Figures S3 B-F**.

Increasing the number of 4-cliques in the network decreases the time to fixation, while not influencing the probability of fixation in regular graphs (**Figure S4**). We only explicitly tune the number of 4-cliques since most other 4-motifs occurrence are correlated with the number of 4-cliques. **Figure S4 B** shows that the effects of the 4-clique on the time to fixation are negligible when the graph contains a low number of triangles and only slightly begins to decrease the time to fixation when the number of triangles in the network is increased. This is surprising since a triangle is a 3-clique and has the opposite effect on time to fixation compared to 4-cliques. However, the observed effects cannot be attributed to 4-cliques alone since increasing the number of 4-cliques can significantly reduce the number of semi-cliques and three-loop-outs, structurally similar to triangles, leading to a net negative influence on the time to fixation. Dependencies between the 4-motifs could explain the observed noise in the time to fixation. Future studies that modify 4 or higher node structures independently could reveal their underlying roles in evolutionary dynamics. Results on higher-order interactions suggest that triangle count is the most effective property in tuning temporal dynamics of a population since the dynamics induced by triangle counts are not affected by other 3-node structures.

### 3 Networks of heterogeneous degree

For networks of heterogeneous degree, node and edge dynamics behave qualitatively similar to those of the *k*-regular graphs, with the dynamics of *Aa* edges faster than those of the nodes. The frequencies of *Aa* edges equilibrate rapidly, as the mutant frequency increases in the population (**Figure S2 A**). These equilibrium *Aa* frequencies decrease as the triangle count increases (**Figure S2 B**). Unlike for regular graphs, the equilibrium *Aa* frequencies decrease by different amounts, with the strongest decreases observed in edges that connect low degree nodes. This difference is also affected by the mixing pattern of the network, with negligible difference in the reduction for graphs with low assortativity. This difference in magnitude causes the probability of fixation to change when the triangle count is altered. Intuitively, if all equilibrium edges are decreased by the same factor, they cancel in the Kolmogorov backward equation, resulting in no net change in the probability of fixation. This is no longer true if the reduction in equilibrium frequencies is not constant across all edge types. As a consequence, the change in probability and time to fixation depend on the equilibrium frequency of *Aa* edges that connect the lowest degree nodes in the graph (**Figure S2 C** and **D**).

Our results generalize to commonly used network families. We first consider small-world networks, with properties observed in many social networks (Watts and Strogatz, 1998). The change in probability of fixation for both death-Birth and Birth-death process due to higher-order interactions is negligible for small and negative assortativity (**Supplementary Figure 5A**). As assortativity increseases, the probability of fixation increases with increasing number of triangles in the network under the Birth-death process (**Supplementary Figure 5B**), and the probability of fixation decreases as triangle counts increase under the dearth-Birth process (**Supplementary Figure 5C**). Times to fixation increase as triangle counts increase for both processes (**Supplementary Figures 5B and D**).

The next graph family we consider is generated using the Barabasi-Albert model of preferential attachment, which creates graphs with the scale-free property often found in social networks (Barabási and Albert, 1999). For this reason, they are typically used to study the spread of information or cultural norms (Cre-anza et al., 2017). The change in probability of fixation for both death-Birth and Birth-death proceses is negligible for these graphs, while the time to fixation increases as triangle counts increase for both processes (**Supplementary Figure 6**).

We also consider random geometric graphs which model spatially structured populations (Waxman, 1988; Penrose et al., 2003). In this family of graphs, nodes have spatial positions randomly drawn from a probability distribution to model spatially homogeneous populations (using the uniform distribution) or populations with heterogeneous spatial density (using the normal distribution). Once the spatial locations of the nodes are determined, the generating algorithm iterates through all pairs of nodes. An edge is created between two nodes if the pair-wise distance is below some prescribed threshold. The change in the probability of fixation for both death-Birth and Birth-death proceses due to higher-order interactions in the network is negligible, while the time to fixation increases as triangle counts increase for both processes (**Supplementary Figure 7**).

---

**Figure S1:**
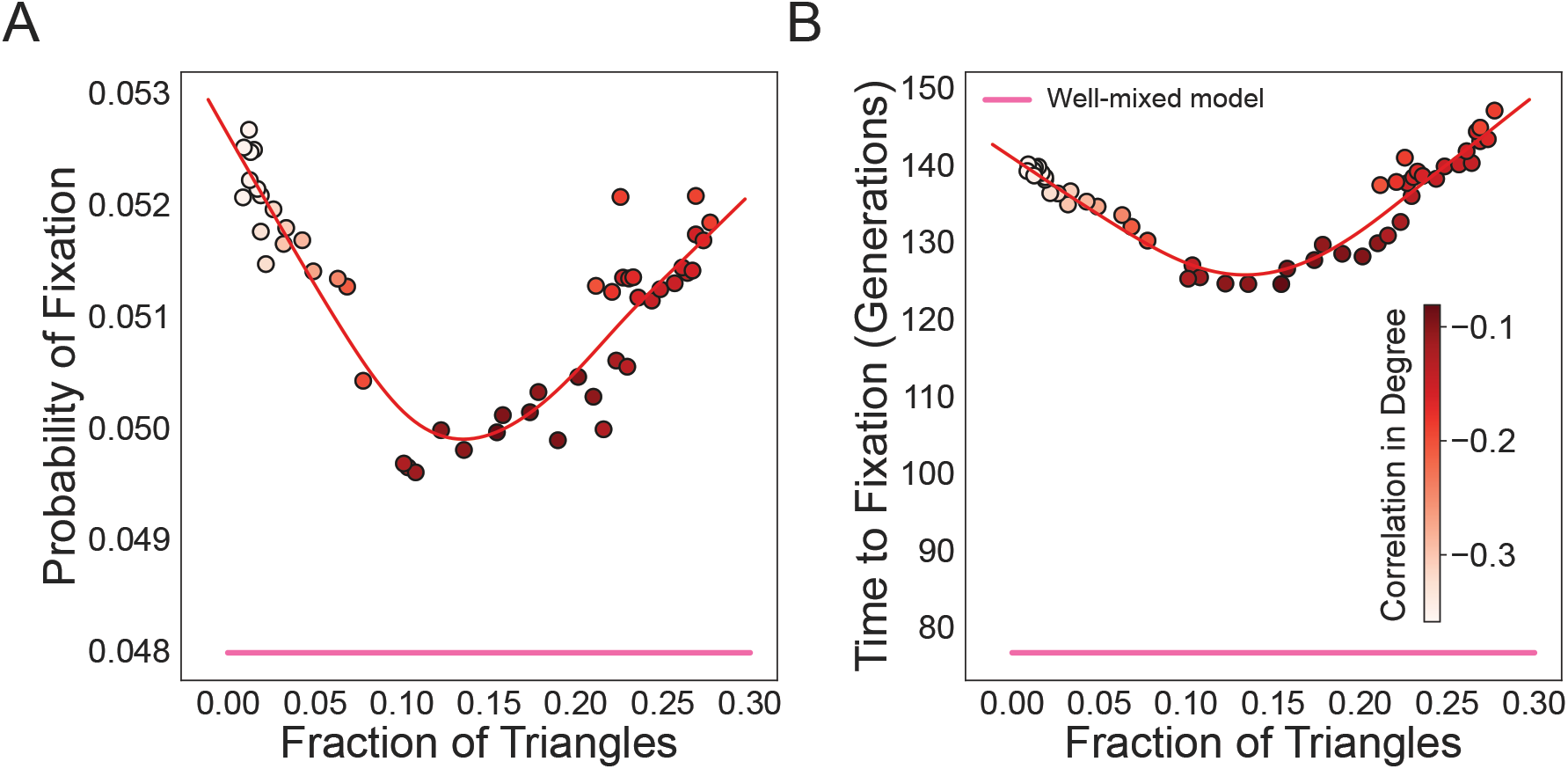
Tuning the fraction of triangles in the network without keeping network mixing pattern constant can result in mistakenly assuming non-linear dependencies. The starting graph is a preferential attachment graph with mean degree equal to five. The degree distribution is held constant. We vary the fraction of triangles in the graph, using edge swapping operations which also modify the mixing pattern of the network, as represented by the color legend. Here, *N* = 100 and *s* = 0.05. The dots represent ensemble averages across 5*e*6 replicate Monte Carlo simulations.

**Figure S2:**
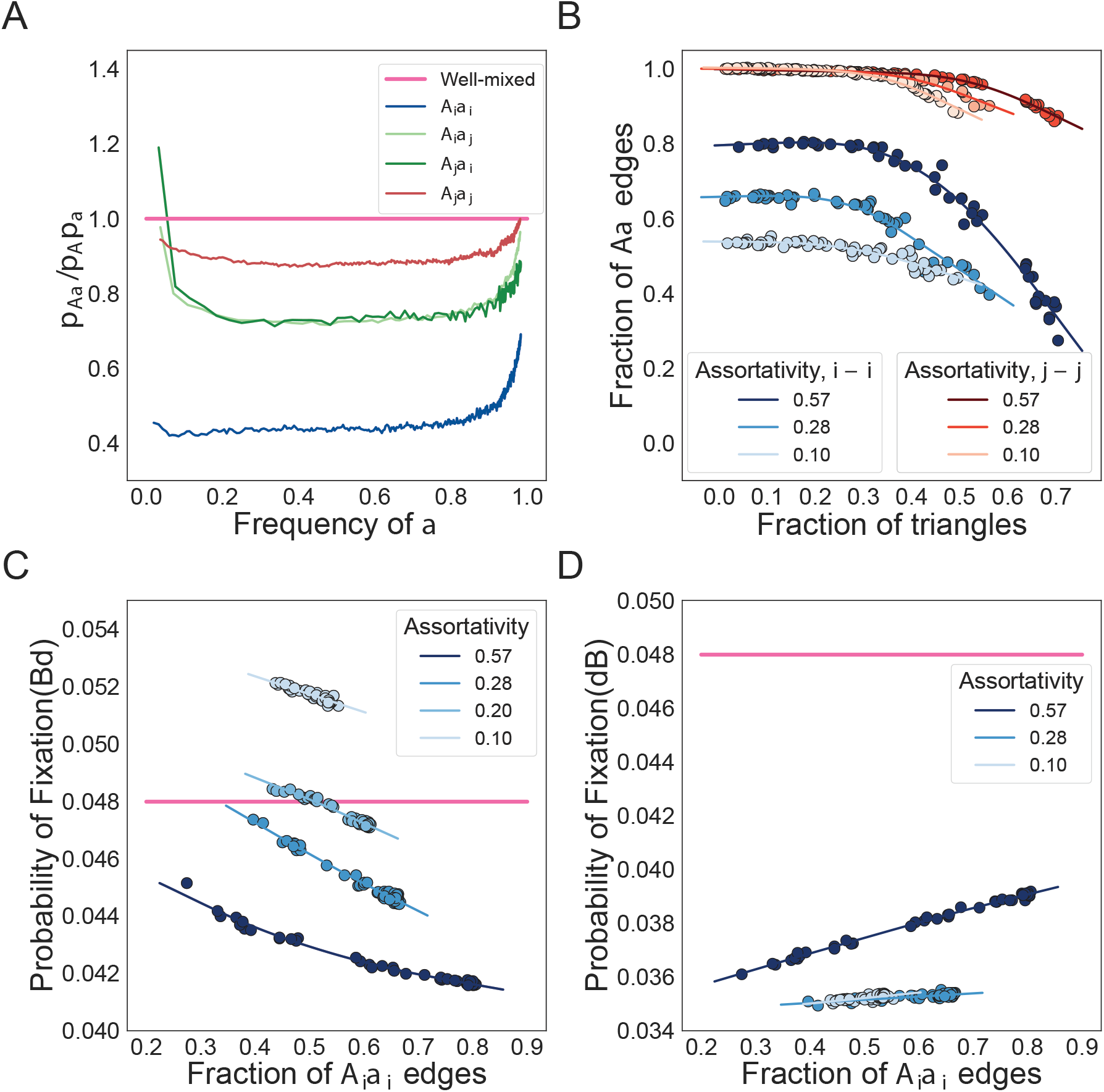
Triangles decrease numbers of different types of *Aa* edges unequally in degree heterogeneous graphs. The degree distribution is held constant as we vary the fraction of triangles in the graphs using edge swapping operations. Here *N* = 100 and *s* = 0.05. Color indicates the degree correlation of the network as a measure of mixing pattern. The dots represent ensemble averages across 5*e*6 replicate Monte Carlo simulations.

**Figure S3:**
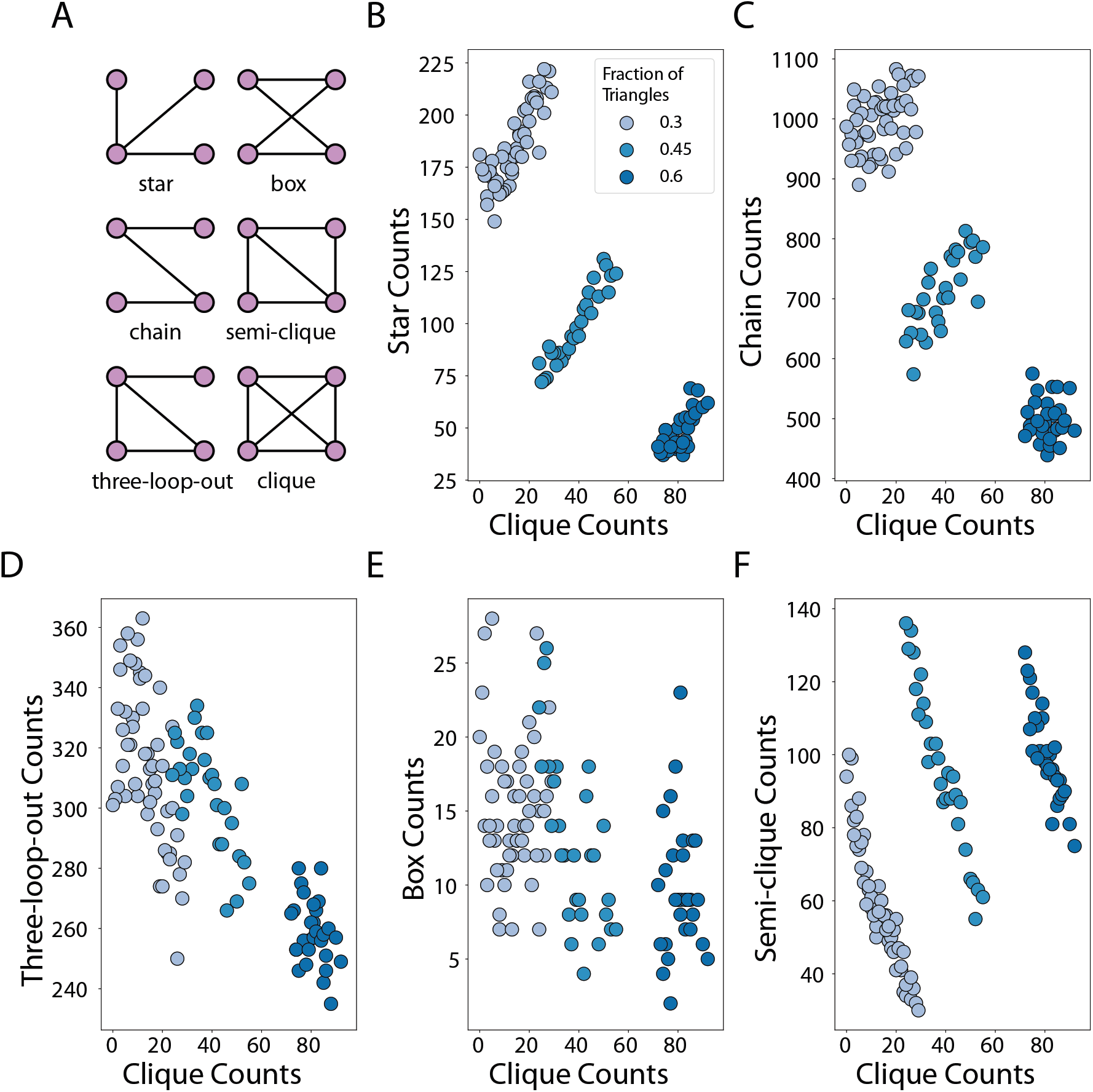
Correlations of 4-motif counts in the graph. We tune the count of 4-cliques in the graph using edge swaps and plot it against the resulting change in the counts of other higher order 4D structures in the graph. **Panel A** showcases the 4-order structures. **Panel B** shows the relationship between the 4-cliques and the stars. **Panel C** shows the relationship between 4-cliques and chains. **Panel D** shows the relationship between 4-cliques and three-loop-outs. **Panel E** shows the relationship between 4-cliques and boxes. **Panel F** shows the relationship between 4-cliques and semi-cliques.

**Figure S4:**
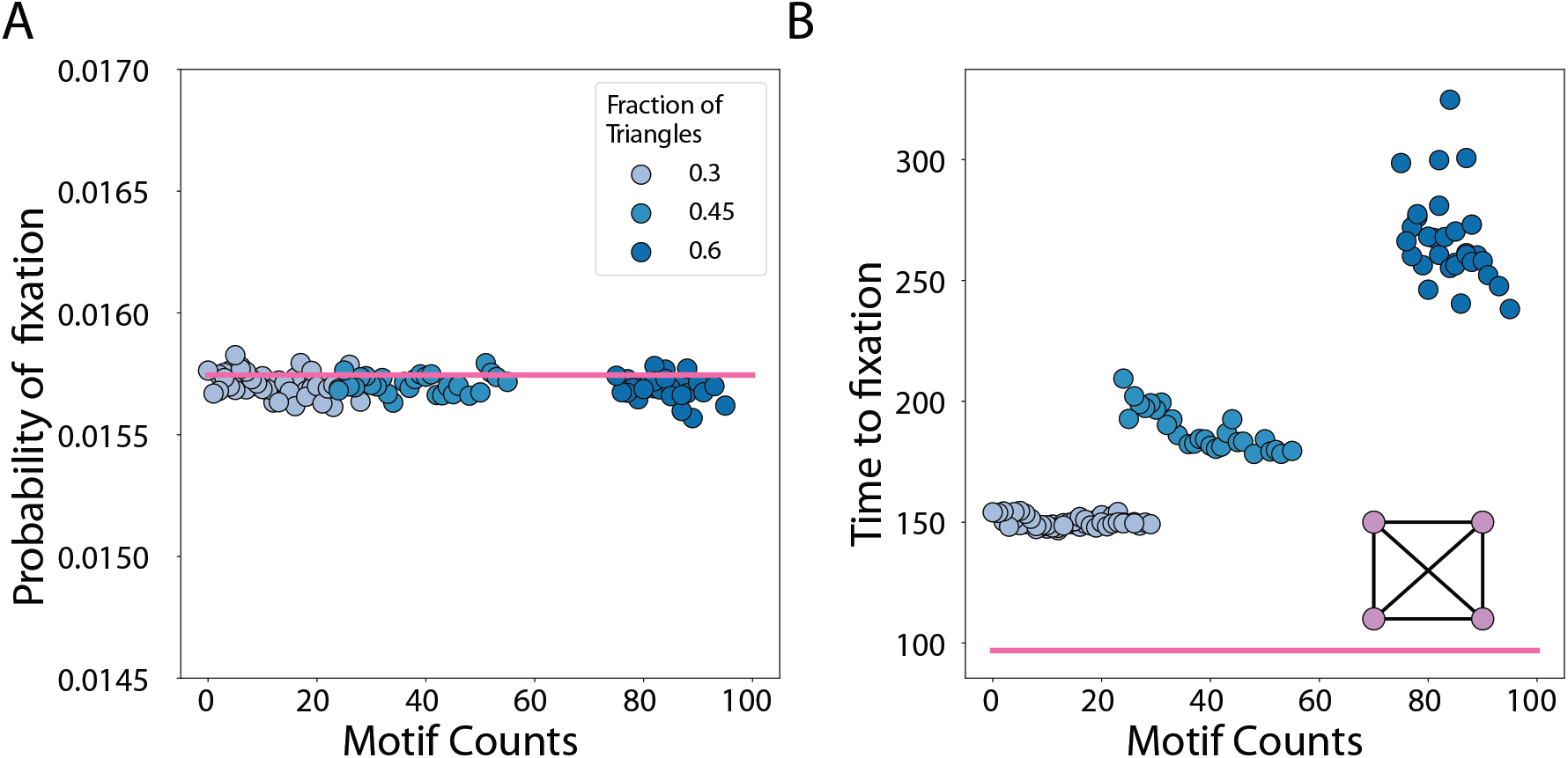
Effects of the fraction of 4-node structures in the graph on evolutionary dynamics. **Panel A**: 4-clique counts in the network do not affect the probability of fixation for regular graphs. **Panel B** Increasing 4-clique counts in the graph decreases fixation time. The dots represent ensemble averages across 5*e*6 replicate Monte Carlo simulations. The colors represent the fraction of triangles in the graph. The graphs used are regular graphs with mean degree of 5 and size of *N* = 100. Here, selection strength *Ns* = 5.

**Figure S5:**
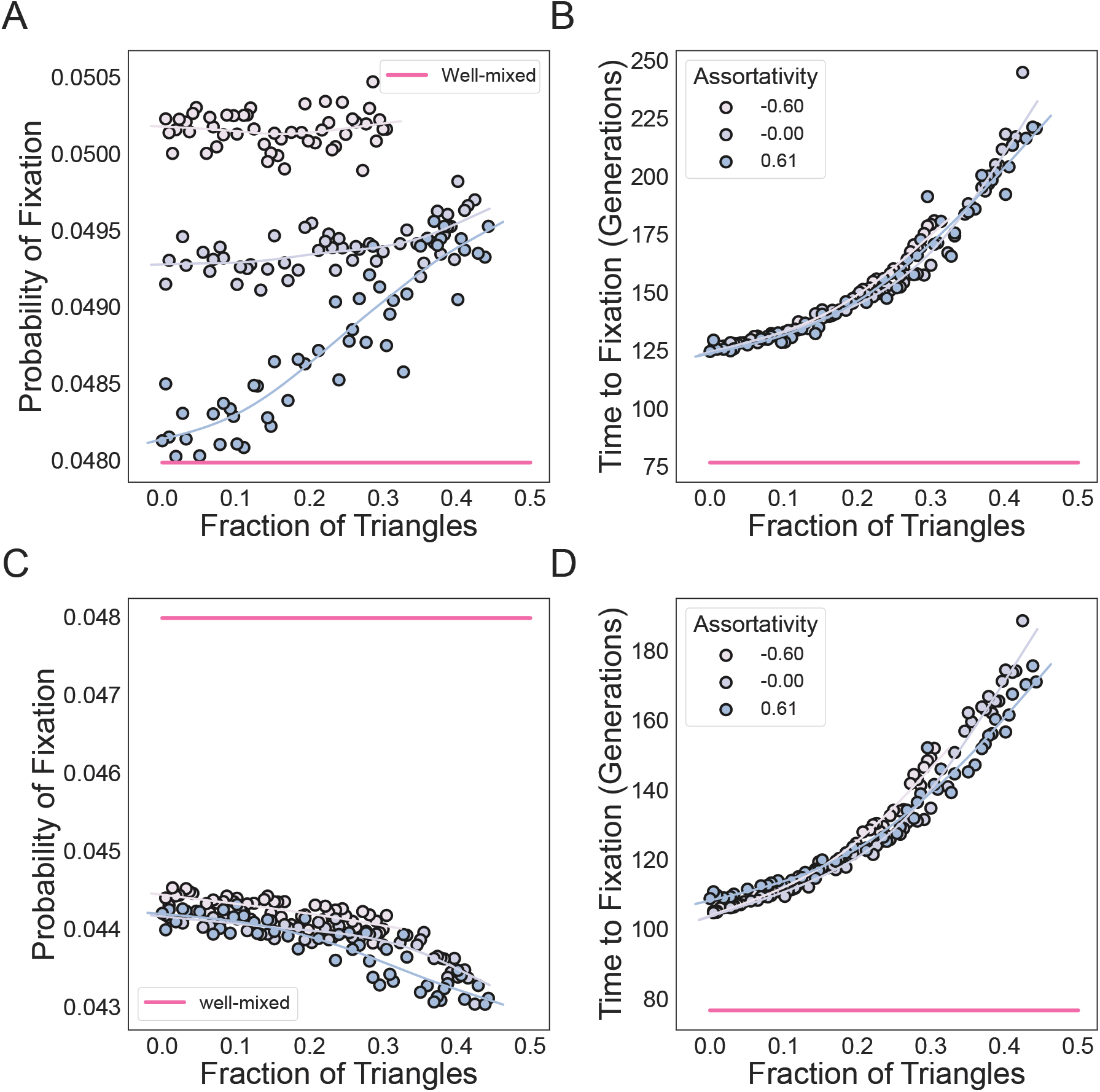
Effects of the fraction of triangle in small-world graphs with various mix patterns. **Panels A** and **B** model the Birth-death process, while **Panels C** and **D** model the death-Birth process. We use small world graphs with mean degree equal to four. The dots represent ensemble averages across 5*e*6 replicate Monte Carlo simulations. The degree distribution and graph assortativity are held constant, as we vary the fraction of triangles in the graphs. The fraction of triangles in the graph is tuned using edge swapping operations. Here *N* = 100 and *s* = 0.05. The colors indicate the assortativity of the network, as in the legend.

**Figure S6:**
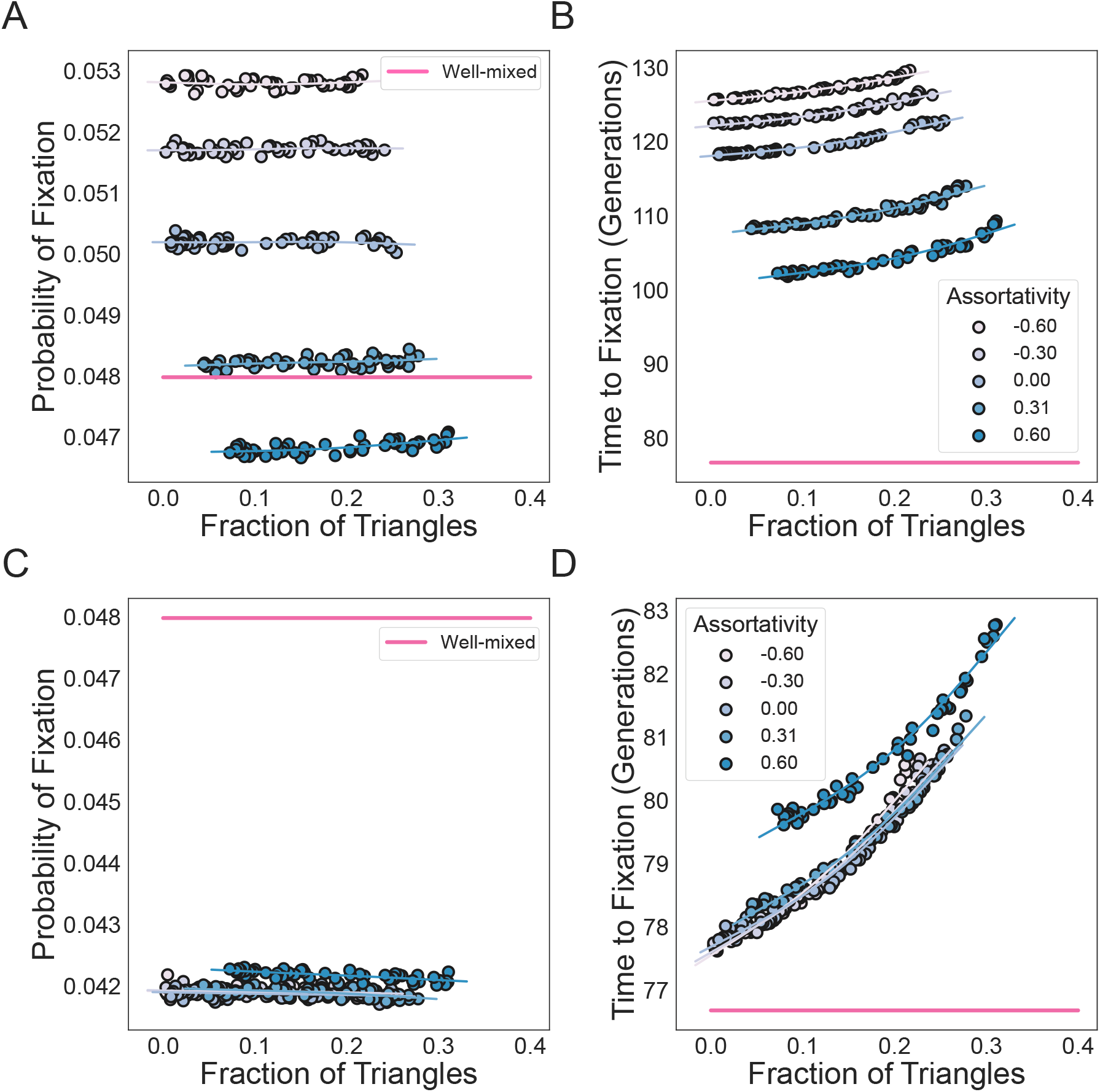
Effects of the fraction of triangle in preferential attachment graphs with various mix patterns. **Panels A** and **B** model the Birth-death process, while **Panels C** and **D** model the death-Birth process. We use preferential attachment graphs with mean degree equal to ten. The dots represent ensemble averages across 5*e*6 replicate Monte Carlo simulations. The degree distribution and graph assortativity are held constant, as we vary the fraction of triangles in the graphs. The fraction of triangles in the graph is tuned using edge swapping operations. Here *N* = 100 and *s* = 0.05. The colors indicate the assortativity of the network, as in the legend.

**Figure S7:**
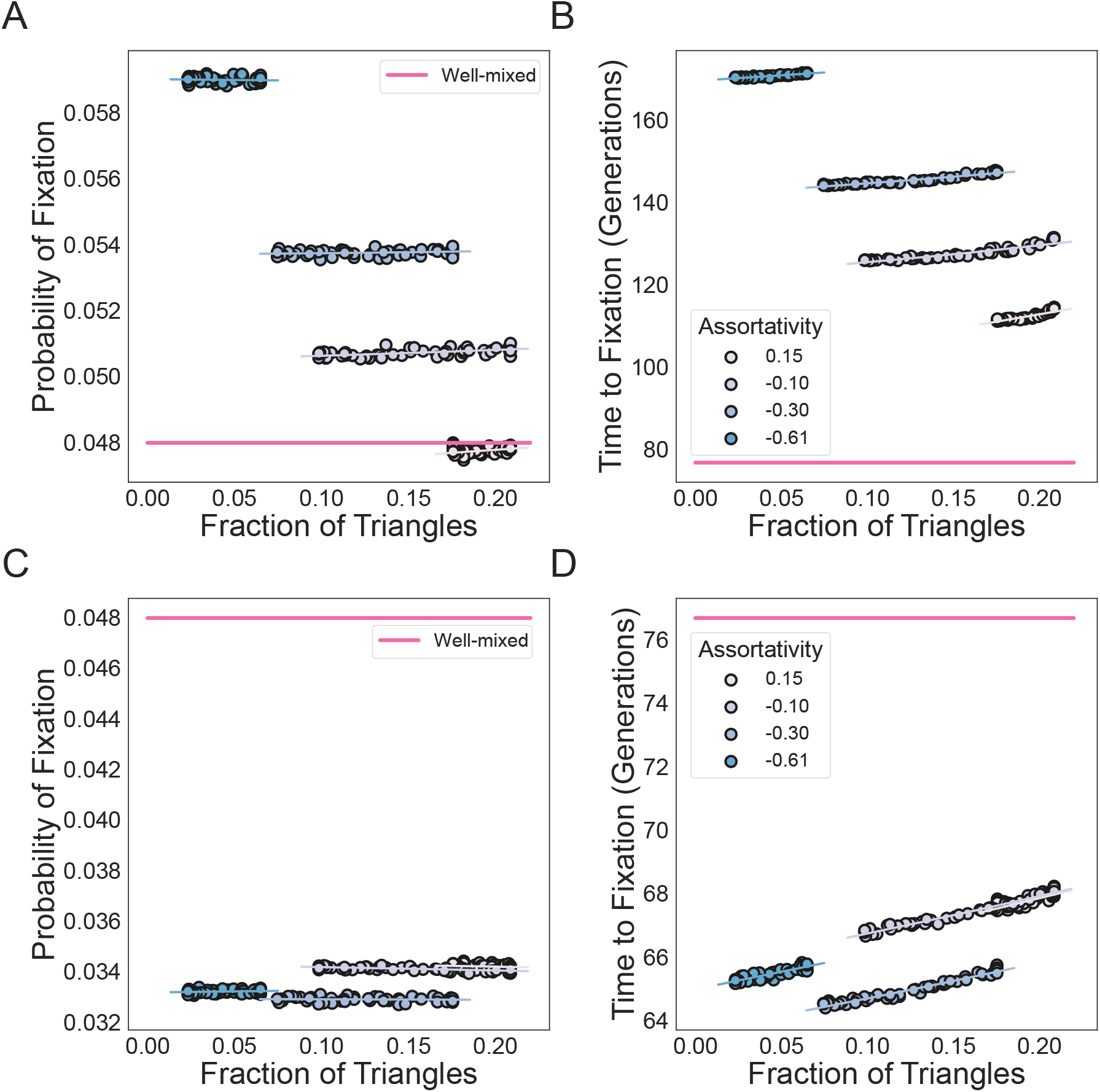
Effects of the fraction of triangle in random geometric graphs with various mix patterns. **Panels A** and **B** model the Birth-death process, while **Panels C** and **D** model the death-Birth process. We use random geometric graphs with a cut-off radius of 0.2. The dots represent ensemble averages across 5*e*6 replicate Monte Carlo simulations. The degree distribution and graph assortativity are held constant, as we vary the fraction of triangles in the graphs. The fraction of triangles in the graph is tuned using edge swapping operations. Here *N* = 100 and *s* = 0.05. The colors indicate the assortativity of the network, as in the legend.

**Figure S8:**
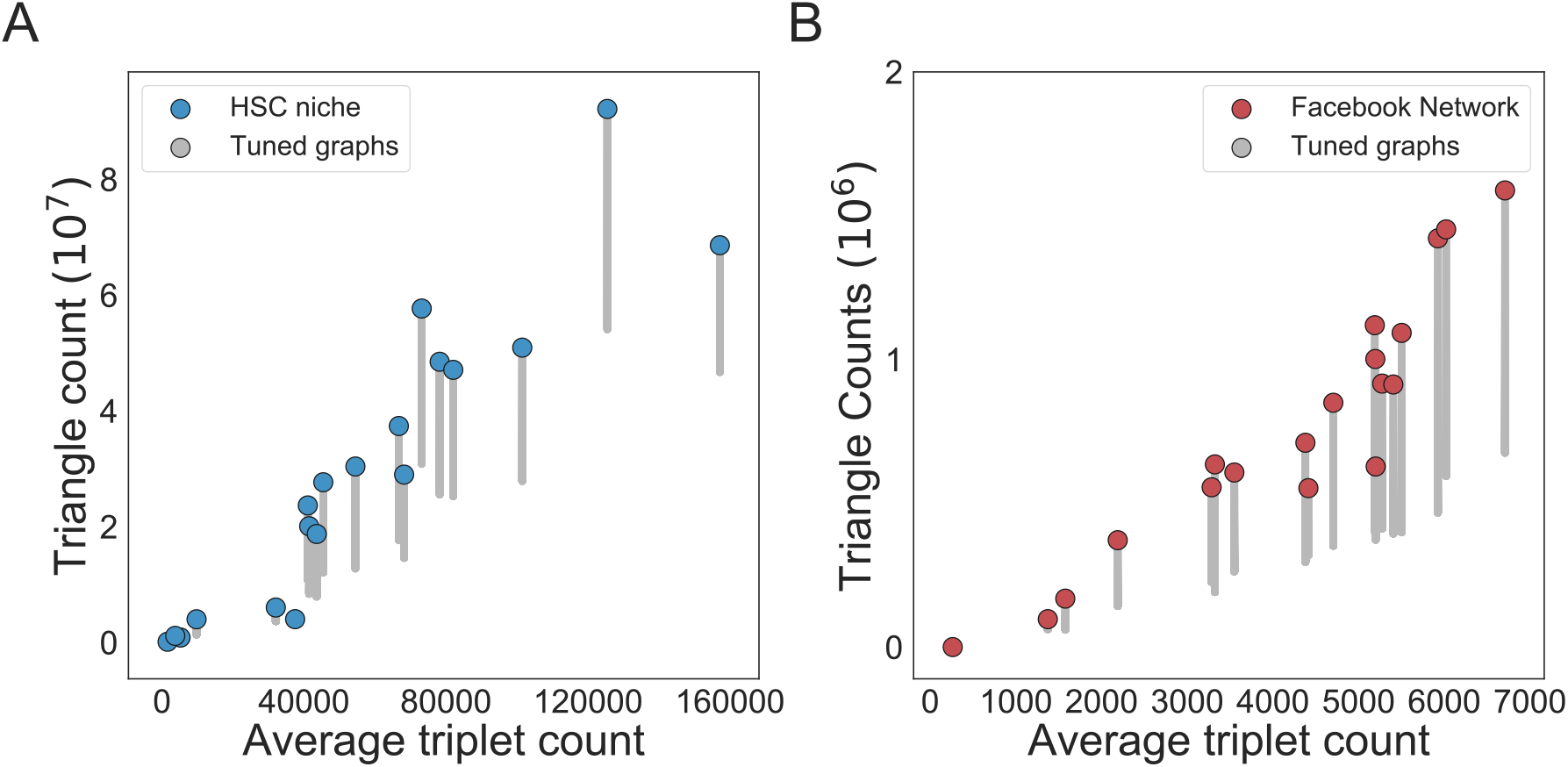
Biological and social populations contain a high number of triangles. Here we take real-world population structures and tune the number of triangles in the network while keeping lower-order structures constant. The triangle count is plotted against the average triplet count possible for those networks. The colored dots represent the original networks and the gray dots represent the tuned networks. In **Panel A** we tune hematopoietic stem cell population structures, while **Panel B** shows social networks, where we tune Facebook friendship networks from across twenty different universities.

## References

1. Philipp M Altrock, Arne Traulsen, and Martin A Nowak. Evolutionary games on cycles with strong selection. Physical Review E, 95(2):022407, 2017.

2. Unai Alvarez-Rodriguez, Federico Battiston, Guilherme Ferraz de Arruda, Yamir Moreno, Matjaž Perc, and Vito Latora. Evolutionary dynamics of higher-order interactions in social networks. Nature Human Behaviour, 5(5):586–595, 2021.

3. Charles W Anderson. Strategy learning with multilayer connectionist representations. In Proceedings of the Fourth International Workshop on Machine Learning, pages 103–114. Elsevier, 1987.

4. Tibor Antal and Istvan Scheuring. Fixation of strategies for an evolutionary game in finite populations. Bulletin of Mathematical Biology, 68(8):1923–1944, 2006.

5. Tibor Antal, Sidney Redner, and Vishal Sood. Evolutionary dynamics on degree-heterogeneous graphs. Physical Review Letters, 96(18):188104, 2006.

6. Marziyeh Askari and Keivan Aghababaei Samani. Analytical calculation of average fixation time in evolutionary graphs. Physical Review E, 92(4):042707, 2015.

7. James Edward Baker. Adaptive selection methods for genetic algorithms. In Proceedings of an International Conference on Genetic Algorithms and their applications, volume 101, page 111. Hillsdale, New Jersey, 1985.

8. Albert-László Barabási and Réka Albert. Emergence of scaling in random networks. Science, 286(5439): 509–512, 1999.

9. Marc Barthélemy. Spatial networks. Physics Reports, 499(1-3):1–101, 2011.

10. Austin R Benson, David F Gleich, and Jure Leskovec. Higher-order organization of complex networks. Science, 353(6295):163–166, 2016.

11. Bryn A Bridges and Roger Woodgate. Mutagenic repair in escherichia coli: products of the reca gene and of the umud and umuc genes act at different steps in uv-induced mutagenesis. Proceedings of the National Academy of Sciences, 82(12):4193–4197, 1985.

12. Mark Broom, Jan Rychtář, and Brian T Stadler. Evolutionary dynamics on graphs-the effect of graph structure and initial placement on mutant spread. Journal of Statistical Theory and Practice, 5(3):369– 381, 2011.

13. Kenneth Mark Bryden, Daniel A Ashlock, Steven Corns, and Stephen J Willson. Graph-based evolutionary algorithms. IEEE Transactions on Evolutionary Computation, 10(5):550–567, 2006.

14. Nicole Creanza, Oren Kolodny, and Marcus W Feldman. Cultural evolutionary theory: How culture evolves and why it matters. Proceedings of the National Academy of Sciences, 114(30):7782–7789, 2017.

15. James F Crow, Motoo Kimura, et al. An introduction to population genetics theory. An introduction to population genetics theory, 1970.

16. Florence Débarre, Christoph Hauert, and Michael Doebeli. Social evolution in structured populations. Nature Communications, 5(1):1–7, 2014.

17. Marcus Frean, Paul B Rainey, and Arne Traulsen. The effect of population structure on the rate of evolution. Proceedings of the Royal Society B: Biological Sciences, 280(1762):20130211, 2013.

17a. Crispin Gardiner. Stochastic methods, volume 4. Springer Berlin, 2009.

18. Philip J Gerrish and Richard E Lenski. The fate of competing beneficial mutations in an asexual population. Genetics, 102:127–144, 1998.

19. Mario Giacobini, Marco Tomassini, Andrea GB Tettamanzi, and Enrique Alba. Selection intensity in cellular evolutionary algorithms for regular lattices. IEEE Transactions on Evolutionary Computation, 9(5):489– 505, 2005.

20. Christos Gkantsidis, Milena Mihail, and Ellen Zegura. The markov chain simulation method for generating connected power law random graphs. Alenex, 2003.

21. Alan Greener, Marie Callahan, and Bruce Jerpseth. An efficient random mutagenesis technique using an e. coli mutator strain. Molecular Biotechnology, 7(2):189–195, 1997.

22. Aric Hagberg, Pieter Swart, and Daniel S Chult. Exploring network structure, dynamics, and function using networkx. Technical report, Los Alamos National Lab.(LANL), Los Alamos, NM (United States), 2008.

23. Mahdi Hajihashemi and Keivan Aghababaei Samani. Fixation time in evolutionary graphs: A mean-field approach. Physical Review E, 99(4):042304, 2019.

24. David Hathcock and Steven H Strogatz. Fitness dependence of the fixation-time distribution for evolutionary dynamics on graphs. Physical Review E, 100(1):012408, 2019.

25. Laura Hindersin and Arne Traulsen. Most undirected random graphs are amplifiers of selection for birth-death dynamics, but suppressors of selection for death-birth dynamics. PLoS Comput Biol, 11(11): e1004437, 2015.

26. Laura Hindersin, Marius Möller, Arne Traulsen, and Benedikt Bauer. Exact numerical calculation of fixation probability and time on graphs. Biosystems, 150:87–91, 2016.

27. Thomas House and Matt J Keeling. Insights from unifying modern approximations to infections on networks. Journal of The Royal Society Interface, 8(54):67–73, 2011.

28. Jeong Han Kim and Van H Vu. Generating random regular graphs. In Proceedings of the thirty-fifth annual ACM symposium on Theory of computing, pages 213–222, 2003.

29. Motoo Kimura. On the probability of fixation of mutant genes in a population. Genetics, 47(6):713, 1962.

30. Yang Ping Kuo, César Nombela Arrieta, and Oana Carja. A theory of evolutionary dynamics on any complex spatial structure. bioRxiv, 2021. doi: 10.1101/2021.02.07.430151. URL https://www.biorxiv.org/content/early/2021/02/08/2021.02.07.430151.

31. Yu-Ping Lai, Jing Huang, Lin-Fa Wang, Jun Li, and Zi-Rong Wu. A new approach to random mutagenesis in vitro. Biotechnology and Bioengineering, 86(6):622–627, 2004.

32. Erez Lieberman, Christoph Hauert, and Martin A Nowak. Evolutionary dynamics on graphs. Nature, 433 (7023):312–316, 2005.

33. Yong Liu, Meng Liang, Yuan Zhou, Yong He, Yihui Hao, Ming Song, Chunshui Yu, Haihong Liu, Zhening Liu, and Tianzi Jiang. Disrupted small-world networks in schizophrenia. Brain, 131(4):945–961, 2008.

34. Priya Mahadevan, Dmitri Krioukov, Kevin Fall, and Amin Vahdat. Systematic topology analysis and generation using degree correlations. ACM SIGCOMM Computer Communication Review, 36(4):135–146, 2006.

35. Elizabeth O McCullum, Berea AR Williams, Jinglei Zhang, and John C Chaput. Random mutagenesis by error-prone pcr. In In vitro mutagenesis protocols, pages 103–109. Springer, 2010.

36. Marius Möller, Laura Hindersin, and Arne Traulsen. Exploring and mapping the universe of evolutionary graphs identifies structural properties affecting fixation probability and time. Communications Biology, 2 (1):1–9, 2019.

37. Jeffrey C Moore and Frances H Arnold. Directed evolution of a para-nitrobenzyl esterase for aqueous-organic solvents. Nature Biotechnology, 14(4):458–467, 1996.

38. Jeffrey C Moore, Hua-Ming Jin, Olga Kuchner, and Frances H Arnold. Strategies for the in vitro evolution of protein function: enzyme evolution by random recombination of improved sequences. Journal of Molecular Biology, 272(3):336–347, 1997.

39. Richard M Myers, Leonard S Lerman, and Tom Maniatis. A general method for saturation mutagenesis of cloned dna fragments. Science, 229(4710):242–247, 1985.

40. Mark EJ Newman. Clustering and preferential attachment in growing networks. Physical Review E, 64(2): 025102, 2001.

41. Mark EJ Newman. The structure and function of complex networks. SIAM review, 45(2):167–256, 2003.

42. Hisashi Ohtsuki, Christoph Hauert, Erez Lieberman, and Martin A Nowak. A simple rule for the evolution of cooperation on graphs and social networks. Nature, 441(7092):502–505, 2006.

43. CJ Paley, SN Taraskin, and SR Elliott. Temporal and dimensional effects in evolutionary graph theory. Physical Review Letters, 98(9):098103, 2007.

44. Andreas Pavlogiannis, Josef Tkadlec, Krishnendu Chatterjee, and Martin A Nowak. Construction of arbitrarily strong amplifiers of natural selection using evolutionary graph theory. Communications Biology, 1 (1):1–8, 2018.

45. Joshua L Payne and Margaret J Eppstein. Evolutionary dynamics on scale-free interaction networks. IEEE Transactions on Evolutionary Computation, 13(4):895–912, 2009.

46. Mathew Penrose et al. Random geometric graphs, volume 5. Oxford university press, 2003.

47. Anatol Rapoport. Spread of information through a population with socio-structural bias: I. assumption of transitivity. The bulletin of Mathematical Biophysics, 15(4):523–533, 1953.

48. L. A. Rastrigin. Systems of extreme control. 1974.

49. Ryan A. Rossi and Nesreen K. Ahmed. The network data repository with interactive graph analytics and visualization. In AAAI, 2015. URL http://networkrepository.com.

50. Vishal Sood and Sidney Redner. Voter model on heterogeneous graphs. Physical Review Letters, 94(17): 178701, 2005.

51. Kenneth O Stanley, Jeff Clune, Joel Lehman, and Risto Miikkulainen. Designing neural networks through neuroevolution. Nature Machine Intelligence, 1(1):24–35, 2019.

52. Angelika Steger and Nicholas C Wormald. Generating random regular graphs quickly. Combinatorics, Probability and Computing, 8(04):377–396, 1999.

53. Felipe Petroski Such, Vashisht Madhavan, Edoardo Conti, Joel Lehman, Kenneth O Stanley, and Jeff Clune. Deep neuroevolution: Genetic algorithms are a competitive alternative for training deep neural networks for reinforcement learning. arXiv preprint arXiv:1712.06567, 2017.

54. Kaustubh Supekar, Vinod Menon, Daniel Rubin, Mark Musen, and Michael D Greicius. Network analysis of intrinsic functional brain connectivity in alzheimer’s disease. PLoS Comput Biol, 4(6):e1000100, 2008.

55. György Szabó and Gabor Fath. Evolutionary games on graphs. Physics Reports, 446(4-6):97–216, 2007.

56. Shaolin Tan and Jinhu Lü. Characterizing the effect of population heterogeneity on evolutionary dynamics on complex networks. Scientific Reports, 4:5034, 2014.

57. Richard Taylor. Contrained switchings in graphs. In Combinatorial Mathematics VIII, pages 314–336. Springer, 1981.

58. Josef Tkadlec, Andreas Pavlogiannis, Krishnendu Chatterjee, and Martin A Nowak. Population structure determines the tradeoff between fixation probability and fixation time. Communications Biology, 2(1): 1–8, 2019.

59. Duncan J Watts and Steven H Strogatz. Collective dynamics of small world networks. Nature, 393(6684): 440–442, 1998.

60. Bernard M Waxman. Routing of multipoint connections. IEEE journal on selected areas in communications, 6(9):1617–1622, 1988.

